# Selective deuteration of an RNA:RNA complex for structural analysis using small-angle scattering

**DOI:** 10.1101/2024.09.09.612093

**Authors:** Aldrex Munsayac, Wellington C. Leite, Jesse B. Hopkins, Ian Hall, Hugh M. O’Neill, Sarah C. Keane

## Abstract

The structures of RNA:RNA complexes regulate many biological processes. Despite their importance, protein-free RNA:RNA complexes represent a tiny fraction of experimentally-determined structures. Here, we describe a joint small-angle X-ray and neutron scattering (SAXS/SANS) approach to structurally interrogate conformational changes in a model RNA:RNA complex. Using SAXS, we measured the solution structures of the individual RNAs in their free state and of the overall RNA:RNA complex. With SANS, we demonstrate, as a proof-of-principle, that isotope labeling and contrast matching (CM) can be combined to probe the bound state structure of an RNA within a selectively deuterated RNA:RNA complex. Furthermore, we show that experimental scattering data can validate and improve predicted AlphaFold 3 RNA:RNA complex structures to reflect its solution structure. Our work demonstrates that *in silico* modeling, SAXS, and CM-SANS can be used in concert to directly analyze conformational changes within RNAs when in complex, enhancing our understanding of RNA structure in functional assemblies.

## INTRODUCTION

RNA:RNA complexes play crucial roles at all levels of gene expression (*1–5*) and can direct various aspects of viral life cycles (*6*). RNAs can fold into a variety of three-dimensional (3D) structures, from simple helices and stem loops, to more complex structures like pseudoknots and triple helices. RNA:RNA complexes form via intermolecular interactions between RNA molecules, either through base pairing or tertiary contacts (*7*, *8*). Knowledge of these complex structures is central to understanding the molecular mechanisms governing RNA function. Despite the importance of RNA structure in directing function, structure elucidation of RNAs and, of particular note, RNA:RNA complexes, has remained relatively unexplored relative to their protein counterparts (*9*). Challenges related to RNA structure determination by high-resolution techniques such as nuclear magnetic resonance (NMR) spectroscopy, X-ray crystallography, and cryogenic electron microscopy (cryo-EM) are best reflected by the paucity of RNA structures in the Protein Data Bank (*10*). Even more sparse are structures of RNA:RNA complexes, which account for a very small fraction of known RNA structures. Thus, there remains a critical need for alternative methods to obtain structural information on RNA:RNA systems.

Small-angle scattering (SAS), with X-rays (SAXS) or neutrons (SANS), is a powerful approach that provides direct insight to RNA structures in solution (*11–14*). In the absence of prior structural information, SAS can rapidly assess the size, shape, oligomerization state, foldedness, and flexibility of an RNA (*15*, *16*). Furthermore, SAS experimental restraints can be combined with computational modeling and molecular dynamics simulations to model tertiary structures (*17–20*). With advancements in the structure prediction of RNAs and their complexes, SAS is a valuable tool for the experimental validation and refinement of predicted atomic models (*21–23*). Additionally, perturbations to the RNA structure, resulting from either changing solution conditions (i.e. temperature or counterion concentration) or complex formation (i.e. binding to a metabolite, protein, or other nucleic acid), are reflected in the scattering data (*16*, *24–26*), making SAS a sensitive reporter of RNA conformation under a wide range of solution conditions.

SANS has a rich history in the structural biology of nucleic acid containing complexes, including the elucidation of the ‘beads-on-a-string’ model of nucleosomes and mapping the relative positions of proteins within the ribosome, long before atomic models became available (*27–29*). In SANS studies, individual components within a complex can be masked via contrast matching (CM), where the scattering length density (SLD, *ρ*) of a specific component is matched to the solvent, such that the resulting signal is dominated by the other component(s) of the complex (*30– 32*). Contrast matching is achieved by manipulating the isotopic composition of the biomacromolecule (via deuterium labeling), the solvent (H_2_O:D_2_O ratio), or a combination of both. The CM-SANS strategy has been successfully applied to elucidate specific structures in protein:protein, protein:nucleic acid, protein:lipid, and nucleic acid:lipid systems, but to date, has not been applied to RNA:RNA complexes (*30*, *32–34*). While theoretical calculations support the feasibility of CM-SANS on selectively deuterated RNA:RNA systems, this approach has not been tested experimentally (*35, 36*).

In this study, we engineered a model RNA:RNA complex to validate our combined SAXS/SANS approach for structure elucidation. First, using SAXS, we structurally characterized the component RNAs individually and in complex. Next, we demonstrated that when one RNA is either perdeuterated (100%) or partially (42%) deuterated, and the other RNA within the complex is protiated, that one RNA component can be contrast matched in an appropriate buffer. To model the RNA:RNA complex, we first modeled the individual RNAs in their bound state by refining an AlphaFold 3 (AF3) predicted complex structure to the experimental SAXS/SANS data. We then compared the quality of the fit [goodness-of-fit (χ^2^)] from the ensemble of bound models to the SANS data. We showed that the models that best fit to the SANS data also best fit to the SAXS data of the complex, improving the AF3-predicted complex and demonstrating that SANS can be used to as a filter to validate RNAs in their bound state. These results highlight the ability to specifically probe the signal of a single RNA within an RNA:RNA complex using SANS to detect subtle binding-induced conformational changes. This hybrid SAXS/SANS approach can be applied to a number of RNA:RNA interactions and therefore has the potential to greatly improve our understanding of quaternary structures of RNA-containing complexes.

## RESULTS

### Design and purification of a model RNA:RNA complex

SAS is an ensemble measurement, therefore it is critical to prepare a sample that remains monodisperse and homogeneous during the time-scale of the experiment (*15*). For the basis of our model, we leveraged a naturally occurring motif from the 5′-leader of the human immunodeficiency virus type 1 (HIV-1) genome, a GC-rich palindromic dimerization initiation sequence (DIS) a key element for genome dimerization, which impacts several different aspects of the HIV-1 life cycle (*37–39*). The DIS is a conserved stem-loop structure, and in isolation, DIS RNAs can form a kissing dimer via intermolecular base pairing between the palindromic loops (5′-GCGCGC-3′ for HIV-1_NL4-3_) with nanomolar affinity (**Fig. 1A**) (*40–42*). To ensure the RNAs form heterodimers with solely intermolecular contacts for SAXS/SANS analysis, we designed two mutant DIS constructs, based on mutations previously used in packaging studies (*43*). In one construct the palindromic loop was mutated to all cytosines (DIS-C) (**Fig. 1A**). In a second construct, the palindromic loop was mutated to all guanines and the hairpin was extended with a UCU bulge and 16 bp elongation (kinked-elongation (*44*), DIS-Gk) to differentiate it from DIS-C (**Fig. 1A**). This design is multipurpose; mutations to the wild-type DIS (DIS-WT) palindrome not only enabled monomeric studies of the DIS-C and DIS-Gk RNAs, but also permitted the differential isotope labeling of the individual components of the DIS-C:DIS-Gk complex for SANS studies. Additionally, as SAS is a low-resolution technique, addition of the kinked elongation motif to DIS-Gk enabled the distinction of the components within the DIS-C:DIS-Gk complex in the surface SAXS envelope.

**Figure 1.**
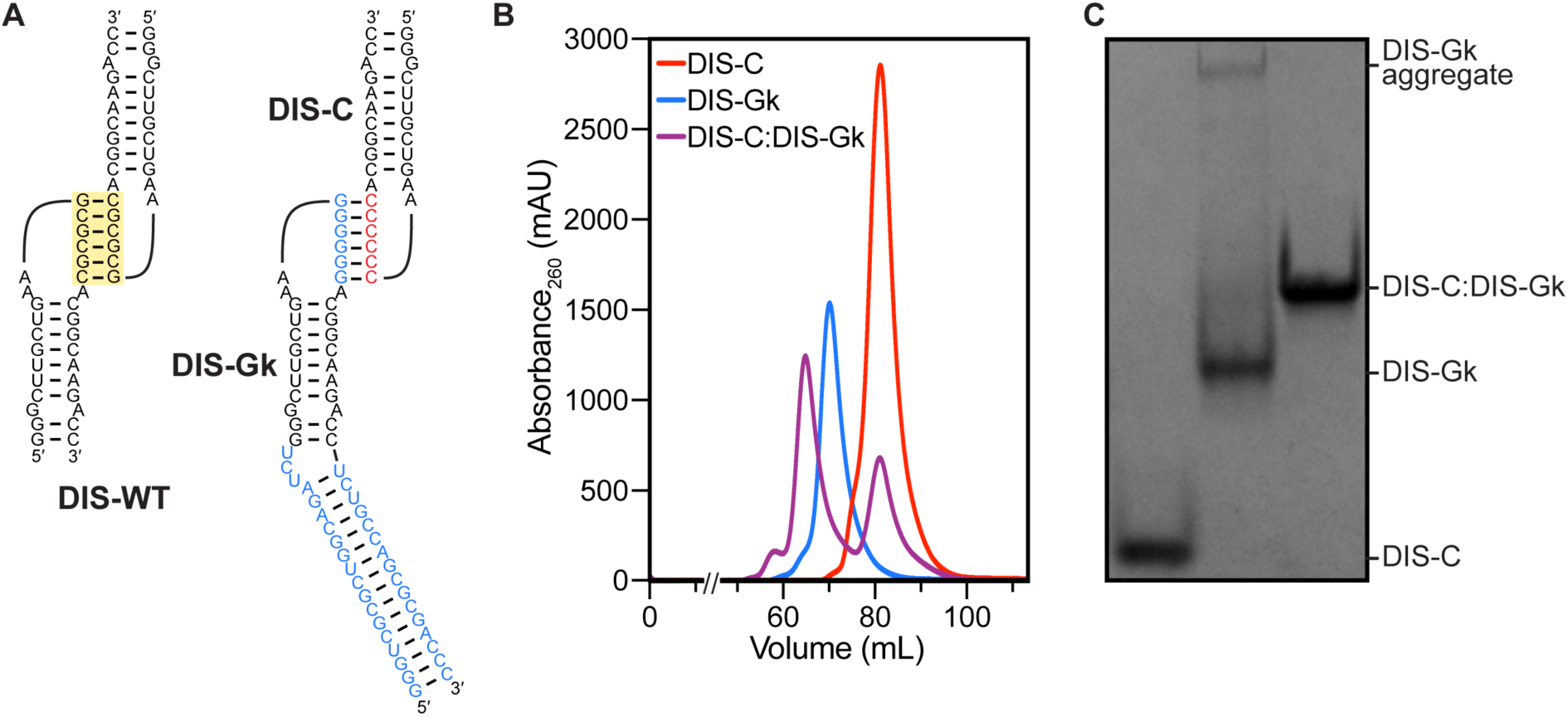
Design and purification of DIS RNAs. **(A)** The DIS-WT palindromic loop (yellow box) was mutated to either all C (red) or all G (blue) to ensure heterodimerization. A kinked-elongation element (blue) was added to DIS-Gk to distinguish it from DIS-C. **(B)** Preparative-scale size-exclusion chromatogram of the DIS RNAs. DIS-C was added in excess during complex formation to saturate DIS-Gk. **(C)** Native gel electrophoresis of the SEC-purified DIS RNAs. After SEC-purification, DIS-C:DIS-Gk complex remains monodisperse.

To ensure the production of a highly homogeneous DIS-C:DIS-Gk complex, we first optimized the preparation of the monomeric hairpin RNAs. Monomeric DIS-C and DIS-Gk RNAs were purified by native size-exclusion chromatography (SEC) (**Fig. 1B**). The purified DIS-C monomer was stable and monodisperse, as monitored by native gel electrophoresis and native SEC-SAXS (**Fig. 1C, Fig. S1A**). However, the DIS-Gk RNA, post SEC-purification, formed a higher-order aggregate in equilibrium, as evidenced by the presence of a slow migrating species on a native gel (**Fig. 1C**). To circumvent issues related to the presence of aggregated DIS-Gk RNA, we used SEC-SAXS, allowing for SAXS data collection of the elution peak corresponding to the DIS-Gk monomer (**Fig. S1B**).

To prepare the DIS-C:DIS-Gk complex, the SEC-purified DIS monomers were incubated together, with excess DIS-C. The DIS-C:DIS-Gk kissing complex was then separated from excess DIS-C via an additional round of SEC purification. Importantly, when the DIS-C:DIS-Gk complex was SEC purified, the complex was monodisperse, with no DIS-Gk aggregate present (**Fig. 1C, Fig. S1C**), allowing for batch mode measurements of the complex using SANS.

### SAXS characterization of DIS-C, DIS-Gk, and the DIS-C:DIS-Gk complex

We first evaluated the global structural and conformational properties of the DIS-C, DIS-Gk, and DIS-C:DIS-Gk RNAs using SAXS. From the SAXS profiles, the Guinier approximation was used to determine the *R_g_* and the intensity at the zero-scattering angle [*I*(*0*)] (**Fig. 2A-C, Table S1D**). The Guinier fit maintains good linearity as *q* approaches zero, indicating good sample quality (**Fig. S2**). Additionally, the Guinier approximation holds true at *qR_g_* ≈ 1.3 for DIS-C and *qR_g_* ≈ 1.0 for DIS-Gk and DIS-C:DIS-Gk, suggesting that DIS-C adopts a spherical shape and DIS-Gk and DIS-C:DIS-Gk adopt more elongated, rod-like shapes (*45*). Guinier analysis yielded *R_g_* values of 14.65 ± 0.04 Å, 26.4 ± 0.1 Å, and 35.3 ± 0.1 Å for DIS-C, DIS-Gk, and DIS-C:DIS-Gk, respectively. The *R_g_* values report on the overall size of the RNAs, with an increasing trend consistent with the design of the respective constructs. From the *I*(*0*), we estimated the molecular weight (MW) via the Porod volume (*V_p_*), yielding MWs of 7.9 kDa, 21.8 kDa, and 30.8 kDa for DIS-C, DIS-Gk, and DIS-C:DIS-Gk, respectively, in agreement with the MWs calculated based on nucleotide sequences (9.3 kDa, 20.8 kDa, and 30.2 kDa, respectively) (**Table S1A,D**).

**Figure 2.**
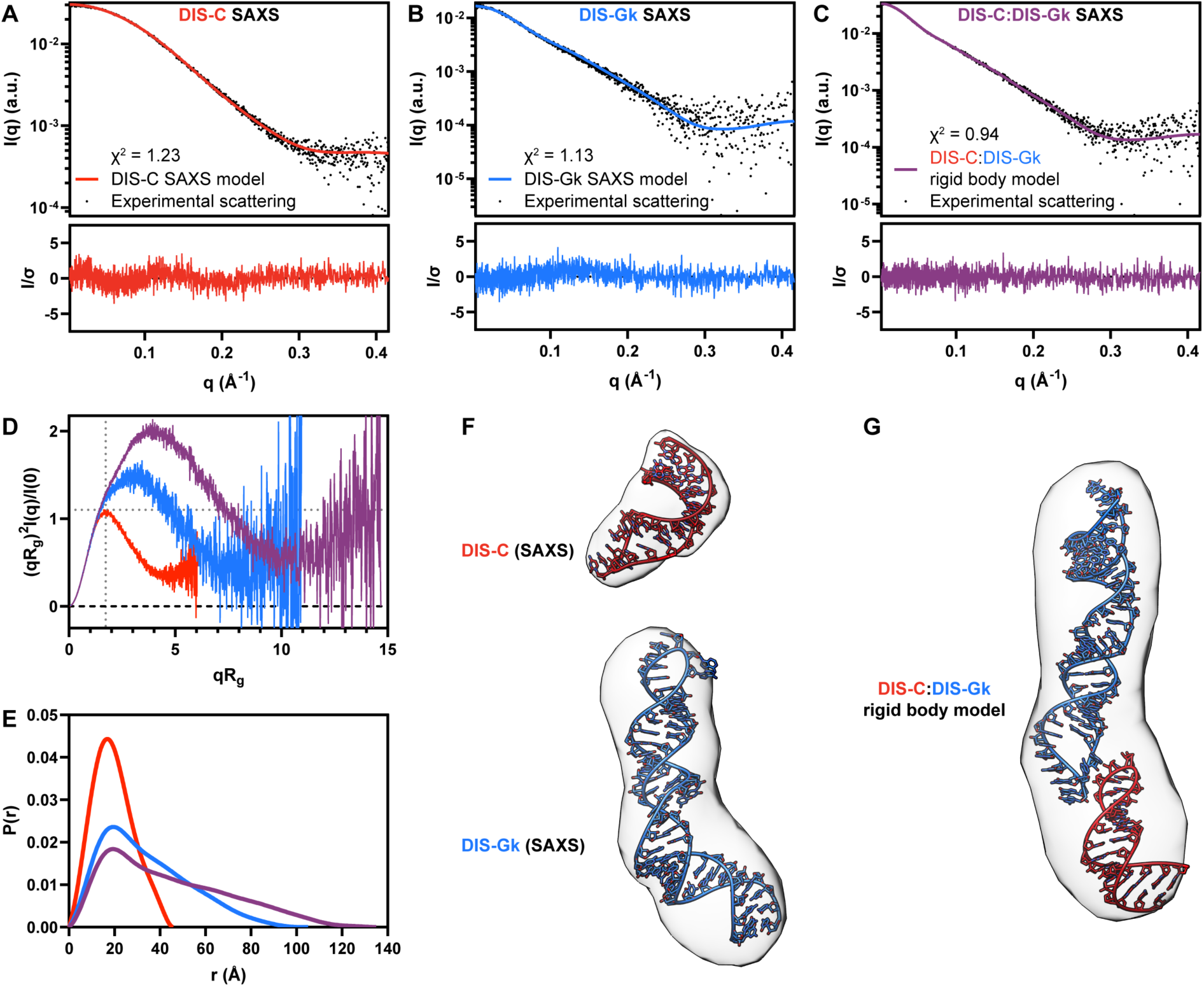
SAXS characterization and modeling of the DIS RNAs. **(A)** DIS-C experimental scattering (black dots) fit to the theoretical scattering of the DIS-C SAXS (red line) model in **(F, top)**. **(B)** DIS-Gk experimental scattering (black dots) fit to the theoretical scattering of the DIS-Gk SAXS (blue line) model in **(F, bottom)**. **(C)** DIS-C:DIS-Gk experimental scattering (black dots) fit to the theoretical scattering of the DIS-C:DIS-Gk rigid body complex (purple line) model in **(G)**. **(D)** Dimensionless Kratky plot of the DIS RNAs. The dashed gray lines on the plot are guidelines for a peak position of a perfectly globular system. **(E)** Normalized pair distance distribution function of the DIS RNAs. **(F)** DIS-C and DIS-Gk structural models using the SAXS data (red and blue, respectively) as restraints. The structural models are fitted to the density map reconstructions (contour levels: 7.5σ). **(G)** SASREF rigid body modeling of structures in **(F)** optimized to the SAXS data in **(C)** fitted to the density map reconstruction (contour level: 7.5σ).

From the *R_g_* and *I*(*0*) parameters derived from the Guinier analysis, we also qualitatively assessed the “foldedness” of the DIS RNAs via a dimensionless Kratky plot (**Fig. 2D**). The DIS-C, DIS-Gk, and DIS-C:DIS-Gk RNAs were all well-folded, with bell-shaped curves (*46*). In agreement with the particle shapes from the Guinier fit, we observe a peak for DIS-C at a (*qR_g_*)^2^*I(q)/I*(0) ≈ 1.1 and *qR_g_* ≈ 1.8, characteristic of a spherical particle with a low degree of flexibility (**Fig. 2D**). However, both DIS-Gk and DIS-C:DIS-Gk deviate from the expected behavior of globular and compacted particles with peaks larger than 1.1 that gradually decrease at high-*q*, consistent with elongated molecules that display a higher degree of flexibility.

For a real-space representation of the DIS RNAs, we also determined the *P*(*r*) function, providing the relative probability associated with sets of distances, *r*, between all pairs of atoms (**Fig. 2E**) (*47*). Further supporting the shape interpretation from Guinier and Kratky analysis, we observed a *P*(*r*) function for DIS-C that resembled a solid sphere, while for DIS-Gk and DIS-C:DIS-Gk, we observed an initial peak with a long extended tail, resembling the *P*(*r*) function of rod-like structures (*48*). All DIS constructs had *P*(*r*) functions that peaked at ∼20 Å, consistent with the presence of A-helical structure. This suggests that in isolation, DIS-C and DIS-Gk exhibit a stem-loop structure, and that the DIS-C:DIS-Gk complex is also mainly helical.

We observed a shoulder in the *P*(*r*) function of DIS-Gk at ∼40 Å, indicative of a bend in the structure, corresponding to the kinked-elongation motif that we introduced. In the DIS-C:DIS-Gk complex, shoulders are apparent at ∼40 Å and ∼60 Å, which likely correspond to the kinked-elongation motif in DIS-Gk and the kissing interaction. Lastly, the *D*_max_ values for DIS-C, DIS-Gk, and DIS-C:DIS-Gk were 46 Å, 105 Å, and 135 Å, respectively, consistent with the increasing size of each sample.

We next performed *ab initio* reconstructions and structure modeling of the DIS RNAs using the SAXS data. *Ab initio* electron density reconstructions using the DENSS algorithm of DIS-C, DIS-Gk, and DIS-C:DIS-Gk RNAs corroborates the overall shapes and structural interpretations from Guinier, Kratky, and *P*(*r*) analysis (**Fig. 2F,G**) (*49*). Furthermore, using RNAMasonry, we modeled 3D structures of DIS-C and DIS-Gk restrained with goodness-of-fits to their respective experimental SAXS data (**Fig. 2F**) (*18*, *50*, *51*). The 3D models of DIS-C and DIS-Gk were fitted to their respective SAXS profiles with χ² values of 1.23 and 1.13, respectively (**Fig. 2A,B**). Both 3D models fit nicely into their respective density map reconstructions (**Fig. 2G**). To determine if the structures reasonably represented the overall shape derived from the DIS-C:DIS-Gk SAXS data, we performed rigid body modeling using SASREF inputting the DIS-C and DIS-Gk models (*52*). Rigid body modeling optimized the monomer structure against the SAXS data, yielding a χ² of 0.94 (**Fig. 2C**). While this confirms that the DIS-C:DIS-Gk reasonably represents the shape of a 1:1 DIS-C:DIS-Gk dimer, it should be noted that the SASREF output flips the orientation of the DIS RNAs in the envelope and positions the RNAs such that intermolecular base pairing and stacking, required for the kissing loop interaction, are prohibited (**Fig. 2G**) (*53*).

Taken together, our SAXS experiments allowed us to characterize the DIS constructs, both in isolation and in complex. The overall complex structure was elongated and the orientation of the DIS-C and DIS-Gk RNAs were discerned due to the presence of the kinked-elongation motif in DIS-Gk. Encouraged by this data, we further characterized these RNAs using contrast variation SANS and isotopic labeling. This approach allowed us to refine the structural models of each individual RNA in the complex and to validate the CM-SANS method.

### Selective deuteration of the DIS-C:DIS-Gk complex

To extract structural information on the DIS-C and DIS-Gk RNAs in their bound states, we turned to CM-SANS. Leveraging the key advantage of SANS over SAXS, where the scattering can be controlled by the isotopic composition rather than the electronic composition of the RNA, we prepared selectively deuterated DIS-C:DIS-Gk complexes (*54*). Deuterium can be incorporated into RNAs enzymatically, via *in vitro* transcription, using commercially available perdeuterated rNTPs, a method routinely used in NMR studies of large RNAs (*55–57*).

Perdeuterated RNAs cannot be contrast matched using H_2_O:D_2_O mixtures, as their scattering length density (7.32 Å^-2^ in 100% D_2_O buffer for a perdeuterated DIS-C) is greater than that of 100% D_2_O solvent (6.36 Å^-2^) (**Fig. 3A**). However, because protiated RNAs can be contrast matched at 65-70% D_2_O (**Fig. 3A**), it is reasonable to make an RNA:RNA complex in which one RNA is perdeuterated and the other RNA is protiated. One drawback from this approach is that while the difference in scattering length density between the perdeuterated and protiated components is maximized, the relatively high content of ^1^H in the contrast matched buffer (35% H_2_O:65% D_2_O) introduces significant background from the incoherent scattering contribution of ^1^H. The incoherent scattering cross section of ^1^H is ∼40 times larger than that of ^2^H (*58*), which decreases the signal-to-noise ratio and presents as high background noise in the scattering data. Therefore, it is ideal to identify the fractional deuteration of an RNA that is contrast matched at a high percent D_2_O buffer to minimized background contributions and to increase the signal-to-noise ratio.

**Figure 3.**
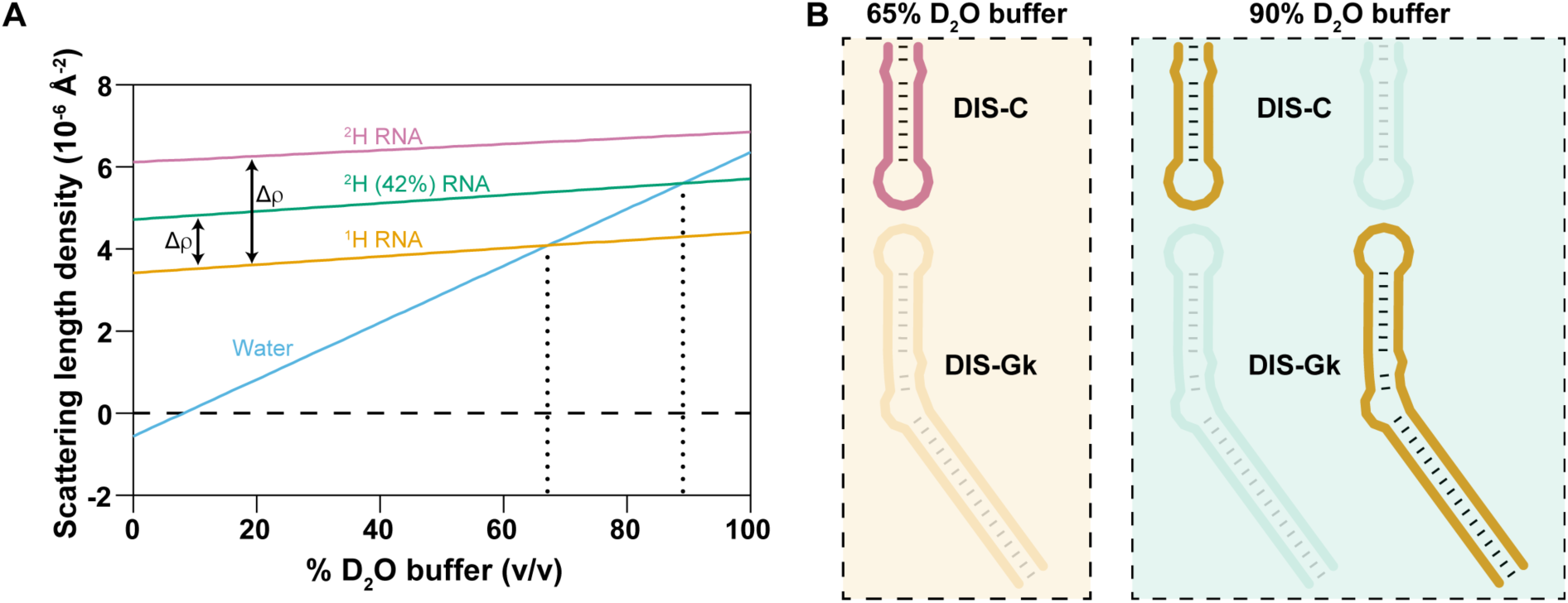
Contrast matching conditions and labeling scheme to probe individual DIS RNAs in complex. **(A)** Calculated scattering length densities (ρ) of protiated RNA (yellow), ^2^H(42%) RNA (teal), and perdeuterated RNA (pink). Contrast between the chosen labeling schemes are denoted as Δρ. The contrast match points (Δρ=0) between the RNAs and appropriate H_2_O:D_2_O buffer are denoted by vertical dotted lines. **(B)** Labeling schemes used in our SANS experiments. The protiated RNA is matched out in 65% D_2_O (left) and the ^2^H (42%) RNAs are matched out in 90% D_2_O (right).

To estimate the contrast match conditions for a partially deuterated RNA, we calculated the theoretical scattering length density for DIS-C, taking into consideration the nucleotide sequence (**Table S1A**) and the number of exchangeable and non-exchangeable hydrogens (**Fig. 3A**). Additionally, we assumed that the excluded volume would remain constant, with only the molecular mass and density increasing after replacement of ^1^H by ^2^H. Random fractional deuteration (42%) of the non-exchangeable hydrogens resulted in a contrast match point of 90% D_2_O buffer for the partially deuterated DIS-C RNA (SLDs of 4.72 Å^-2^ and 5.71 Å^-2^ in 0% and 100% D_2_O buffer, respectively). We employed the same calculations and deuteration strategy for DIS-Gk (**Table S1A**). As a control, we experimentally validated the contrast match point of ^2^H(42%)-DIS-C to be at 90% D_2_O buffer (**Fig. S3**). This result agrees with previous findings where the deuteration of 50S subunit ribosomal RNA from *E. coli* cells grown in 65% D_2_O incorporated 31% ^2^H, resulting in a contrast match point of approximately 80% D_2_O (*59*).

We chose two experimental buffer conditions: 65% D_2_O and 90% D_2_O, within the predicted contrast match points of protiated RNA and 42% deuterated RNA, respectively (**Fig. 3A**). Our labeling strategy consisted of three selectively deuterated complexes: ^2^H-DIS-C:^1^H-DIS- Gk and ^1^H-DIS-C:^2^H(42%)-DIS-Gk, both probing the bound DIS-C, and ^2^H(42%)-DIS-C:^1^H-DIS- Gk, probing bound DIS-Gk (**Fig. 3B**).

### SANS characterization of bound DIS-C and bound DIS-Gk in complex

To validate that individual RNAs within an RNA:RNA complex could be contrast matched, we performed SANS experiments on a selectively deuterated DIS-C:DIS-Gk complex. As a proof-of-principle, our first SANS experiment was conducted on the ^2^H-DIS-C:^1^H-DIS-Gk complex in 65% D_2_O, masking out the protiated DIS-Gk (**Fig. 4A, Fig. S4A**). Despite this sample showing signs of subtle aggregation in the low-*q* region, Guinier derived *R_g_* of ^2^H-DIS-C:^1^H-DIS-Gk complex yielded an *R_g_* of 13.3 ± 1.8 Å (**Fig. S5A**, **Table S1D**), suggesting that the much larger DIS-Gk is contrast matched to the solvent and that the main contributions to scattering come from DIS-C within the complex.

**Figure 4.**
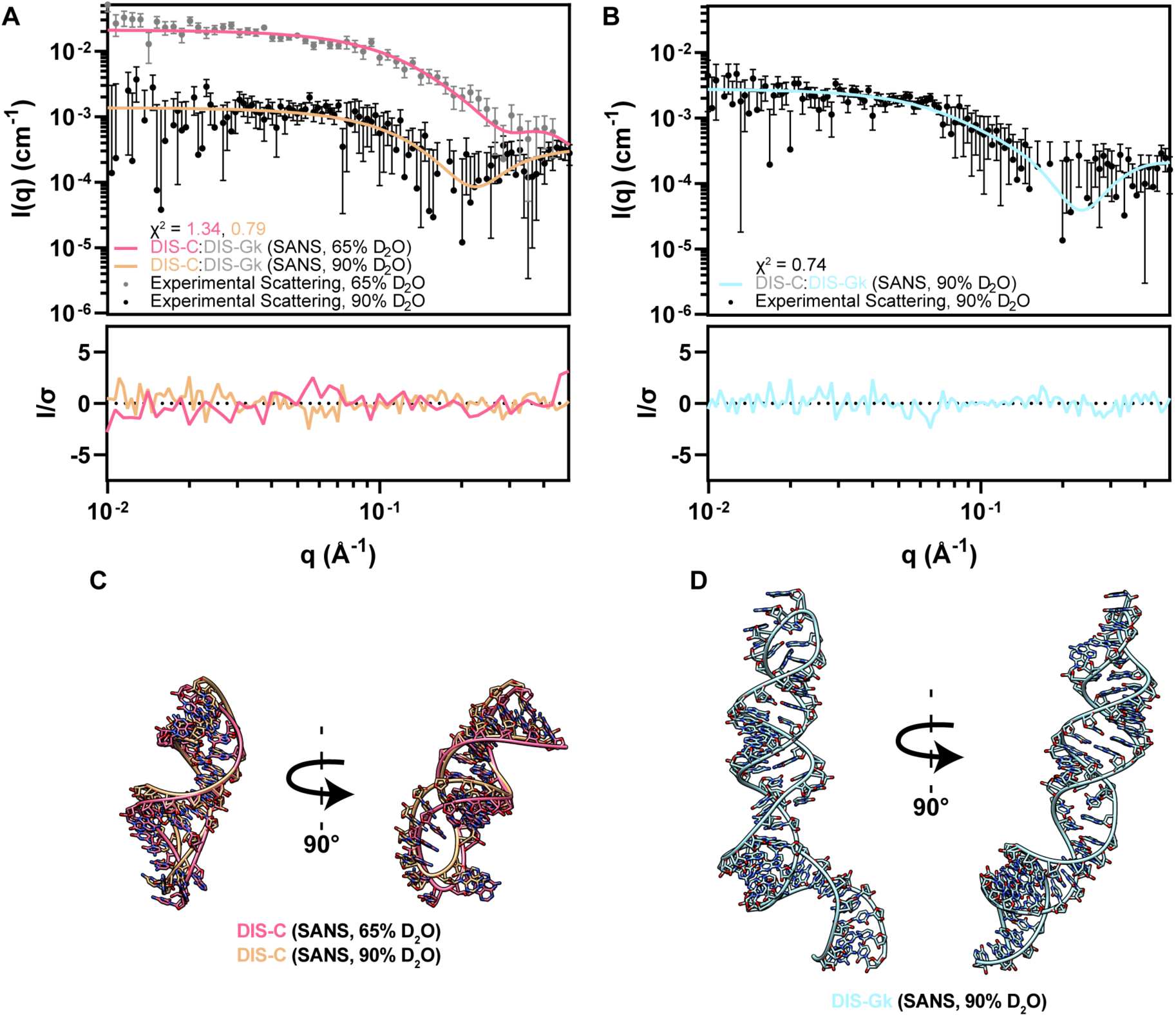
CM-SANS allows for modeling and validation of the DIS RNAs in their bound state. **(A)** Experimental scattering of ^2^H-DIS-C:^1^H-DIS-Gk in 65% D_2_O (gray dots) and ^1^H-DIS-C:^2^H(42%)-DIS-Gk in 90% D_2_O (black dots), where DIS-Gk is matched out. Theoretical scattering profiles of DIS-C structural models in **(C)** are fitted against their respective experimental SANS data. **(B)** Experimental scattering of ^2^H(42%)-DIS-C:^1^H-DIS-Gk in 90% D_2_O (black dots), where DIS-C is matched out. Theoretical scattering profiles of the DIS-Gk structural model in **(D)** is fitted against the experimental SANS data. **(C)** DIS-C structural models using the SANS data as restraints [pink (65% D_2_O) and light orange (90% D_2_O)]. **(D)** DIS-Gk structural model using the SANS data as restraints [light teal (90% D_2_O)]. *q*-points without downward error bars in **(A)** and **(B)** have a non-positive lower bound, which cannot be shown on a log axis.

We next performed SANS experiments on ^1^H-DIS-C:^2^H(42%)-DIS-Gk and ^2^H(42%)-DIS-C:^1^H-DIS-Gk in 90% D_2_O. This condition minimized the scattering contribution from the partially deuterated component and maximized scattering from its protiated counterpart while providing an additional contrast point for DIS-C (**Fig. 3B**). The experimental SANS profiles under different contrast conditions (**Fig. 4A,B, Fig. S4A,B**) were consistent with our simulated SANS profiles (**Fig. S6**). We determined Guinier-derived *R_g_* values of 14.1 ± 3.5 Å and 26.7 ± 3.3 Å for DIS-C and DIS-Gk, respectively, in the complex, suggesting that there are no scattering contributions from the matched-out component (**Fig. S5B,C, Table S1D**). Additionally, we determined the *P*(*r*) functions for the selectively deuterated complexes. For ^2^H-DIS-C:^1^H-DIS-Gk in 65% D_2_O and ^1^H-DIS-C:^2^H(42%)-DIS-Gk in 90% D_2_O, which probed DIS-C in complex, the *D*_max_ values were 53 Å and 46 Å, respectively (**Fig. S7A, Table S1D**). For ^2^H(42%)-DIS-C:^1^H-DIS-Gk in 90% D_2_O, which probed DIS-Gk in complex, the *D*_max_ value was 97 Å (**Fig. S7B, Table S1D**). These SANS-derived *D*_max_ values align with the SAXS-derived *D*_max_ values of free DIS-C (46 Å) and DIS-Gk (105 Å) and was less than that of the DIS-C:DIS-Gk complex (135 Å), showing effective contrast matching of one RNA in the complex (**Fig. S7, Table S1D**). We modeled 3D structures of DIS-C and DIS-Gk using RNAMasonry, restrained with goodness-of-fits to their respective experimental SANS data (**Fig. 4C,D**) (*18*). We generated two models for DIS-C, fitted to the data collected at 65% D_2_O (χ^2^ = 1.34) and 90% D_2_O (χ^2^ = 0.79), and one model for DIS-Gk, fitted to the data collected at 90% D_2_O (χ^2^ = 0.74) (**Fig. 4A,B**). The theoretical scattering profiles agree well with the experimental scattering, further supporting that the SANS data reflect structural features of the specifically probed RNA.

To determine if these SANS-derived models were consistent with the isolated RNAs, we fitted them to their respective SAXS data. Both DIS-C SANS models have suboptimal fits to the monomeric DIS-C SAXS data [χ² = 2.05 (65%) and χ² = 3.57 (90%)], suggesting that there are conformational changes in DIS-C when bound (**Fig. S8, Table S2**). Comparably, the DIS-Gk SANS model has a reasonable but suboptimal fit to the monomeric DIS-Gk SAXS data (χ² = 1.88), suggesting that the perturbations of the bound state are quite small relative to the global ensemble captured in solution (**Fig. S9, Table S3**). Rigid body modeling using the SANS-derived structural models represent the overall shape of the complex well (**Fig. S10**). However, as with the SAXS complex rigid body model, the SASREF output has the incorrect orientation and does not include reasonable base pairing or stacking characteristic of RNA:RNA kissing interactions. This underscores a limitation in SAXS-only comparisons of complexes, where multiple conformations or orientations of the individual components can satisfy the scattering of the overall complex.

### Modeling of the DIS-C:DIS-Gk kissing complex to the SAS data

To model a physical complex with proper base pairing interactions, we used AF3 (*21*) to generate starting models of the DIS-C:DIS-Gk complex (**Fig. S11A**). These AF3 models were poor fits to the DIS-C:DIS-Gk complex SAXS data, with a χ² range of 3.66 to 6.22 (**Fig. S11B, Table S4**), indicating that they don’t reflect the true conformation of the complex in solution. Interestingly, while the AF3 models were poor fits to the SAXS scattering data, the apical loops in these models adopted structures consistent with a kissing interaction. We next generated chimeric structural models in which the apical loops from the AF3 models (with the kissing loop stacking geometry) were appended to the stems of the DIS-C and DIS-Gk RNAMasonry models (**Fig. S12A-C**). These DIS-C and DIS-Gk structures, which include the AF3 kissing loops, were further refined against either our SAXS or SANS data. During refinement, the kissing loop residues were excluded from the coarse-grained modeling (**Fig. 5A, Fig. S12D**). To identify which of the refined models best represented the bound state of DIS-C and DIS-Gk, we generated theoretical scattering profiles for each model and compared them to our experimental SANS data. Overall, the DIS-C + KL (SANS, 65% D_2_O) had the best fit to the SANS data collected in 65% D_2_O (χ² = 1.22) and had a roughly similar fit to the other models against the 90% D_2_O (χ² = 0.80) SANS data (**Figs. S13, S14, Table S2**). Interestingly, the DIS-Gk + KL (SAXS) model (χ² = 0.74) better fit the SANS data (**Fig. S15, Table S3**).

**Figure 5.**
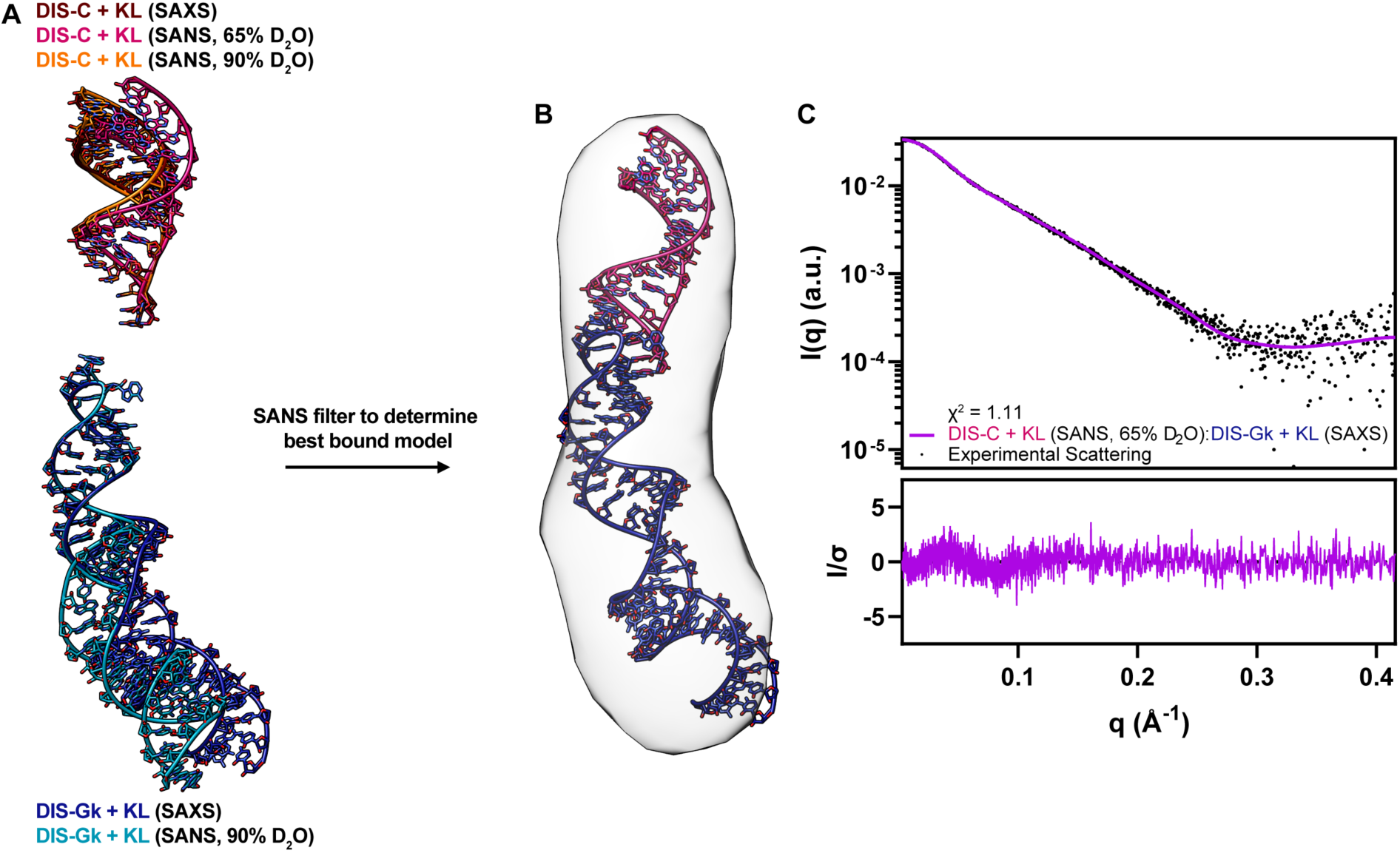
SANS validation of the kissing loop models provides the best reasonable complex model. **(A)** DIS-C and DIS-Gk models restrained to the SAS data while maintaining proper kissing loop geometry from the AF3 models. **(B)** Back-calculation of the individual models in **(A)** to their respective SANS data determined the best models that represented the bound state. **(C)** Theoretical scattering of the best model, DIS-C + KL (SANS, 90% D_2_O):DIS-Gk + KL (SAXS), fit to the DIS-C:DIS-Gk SAXS data with corresponding residuals.

Using the best models as judged by the SANS fits, we aligned the kissing loop residues of DIS-C + KL (SANS, 65% D_2_O) and DIS-Gk + KL (SAXS) to the kissing loop residues of their respective RNAs in the AF3 predicted DIS-C:DIS-Gk complex (**Fig. 5B**) and back-calculated the theoretical scattering of the complex. The DIS-C + KL (SANS, 65% D_2_O):DIS-Gk + KL (SAXS) complex was a vast improvement relative to the initial AF3 models, fitting the DIS-C:DIS-Gk complex SAXS data with a χ² = 1.11 (**Fig. 5C**). Notably, a relationship can be drawn when comparing all six possible combinations of the DIS + KL complex from our DIS + KL models. Generally, complexes generated with DIS-C + KL and DIS-Gk + KL models that best represent their bound structures (models with the lower χ² values relative to the respective SANS data) also had lower χ² values against the complex SAXS data, underscoring the utility of SANS to detect the representative bound state (**Fig. S16, Table S5**). It should be noted though that due to the large error bars for our 90% D_2_O SANS data, the statistical power of the χ² test is reduced, leading to a lower absolute χ² value. However, a relative comparison can still be made when comparing models to the same dataset by looking at their relative difference, where the effect of the noise is consistent across models.

Although the initial manual alignment of SAXS/SANS-guided models to the AF3 predicted complex worked well to a first approximation, we wanted to optimize the complex structure. To probe the potential dynamics of the complex, we used normal mode analysis to estimate the flexibility of the DIS-C + KL (SANS, 90% D_2_O):DIS-Gk + KL (SAXS) complex and further improve the agreement to the SAXS data using SREFLEX (*60*). The best conformer showed a marginal improvement, with a χ² = 1.07 and overall RMSD of 4.1 Å, opening a few base pairs in the complex (**Fig. S17**). Given the inherent dynamic nature of RNAs, and the DIS RNA specifically (*61–63*), the modeled flexibility may reflect the behavior of the experimental ensemble. This dynamic behavior, however, is not fully captured in the initial structural modeling, which is constrained by stringent base-pairing requirements. Nonetheless, the correspondence between our top SAXS/SANS-guided model and the experimental data suggests that the model provided a reasonable approximation of the experimental observations. Future algorithms capable of directly modeling RNA:RNA complexes using SAS experimental restraints could significantly enhance the optimization and streamlining of complex structural determinations. However, substantial advancements in computational methods and integration of experimental data are still required to achieve this goal (*64*, *65*).

Overall, based on the SANS data, we demonstrated that there are differences between the SANS structural ensemble exhibited by the DIS RNAs in complex relative to the SAXS structural ensemble of the DIS RNAs in isolation. Additionally, using the SANS data as a validation and refinement tool, we determined the models that best represented the bound state. These models showed an improved fit to the experimental SAXS data of the complex, highlighting the sensitivity and utility of a combined SAXS/SANS approach to elucidate conformational rearrangements of RNA:RNA complexes.

## DISCUSSION

Computational and biophysical methods that provide insight into the structure and dynamics of RNA:RNA interactions are of great interest. While methods to determine the secondary and tertiary structures of RNAs are in constant development, characterization of RNA:RNA complexes still lags behind other biomolecular complexes (*64*, *66*, *67*). SAXS and SANS are valuable tools in integrative structural studies of biological complexes (*30*, *33*, *68*). However, applications on RNA:RNA complexes have been few, often combining SAXS with other techniques to inform only on the overall complex structure (*25*, *67*, *69*).

Our study is the first to investigate structural perturbations of RNA-only systems upon complex formation using a joint SAXS/SANS approach by leveraging selective deuteration and contrast matching. Traditionally, SANS studies of RNA:RNA complexes have been hampered by difficulties in preparing deuterated RNAs (*33*), but advancements in RNA preparation have made this more accessible (*55*, *57*, *70*). As an alternative approach, contrast variation SAXS, offers higher sensitivity and has been applied to the study of protein:nucleic acid complexes (*71*). However, to achieve contrast matching, of an RNA:RNA complex, the electronic composition of one of the component RNAs would need to be changed. While it is possible to chemically modify the sugar-phosphate backbone (*72*), this approach is less practical than isotope labeling as it can alter the structure of the RNA (*73*, *74*). Consequently, SANS remains an appealing choice due to ease in isotope labeling and the capability of performing experiments under physiologically relevant conditions.

While our approach depended on the hybridization of two RNAs that were independently labeled, segmental labeling approaches can be used to change the isotope composition of single biomolecules for structural studies (*75–82*). In fact, through segmental labeling of an alternative splicing factor, SANS could be used to define the domain arrangement of the multi-domain protein (*24*). Advances in the segmental labeling of RNAs (*82*) opens the door for similar studies of large, multi-domain RNAs. Our work, which demonstrated the feasibility of contrast matching selectively deuterated RNAs within and RNA:RNA complex, lays the foundation for the study of larger RNA:RNA or RNA:protein complexes where segmental labeling approaches could be applied to allow for the direct observation of specific domains within a large transcript.

While advancements in structure prediction algorithms such as AF3 have allowed for the prediction of RNA:RNA complexes from sequence alone, the accuracy and confidence of the models are still far from those of proteins (*21*). Thus, comparison with experimental data, such as SAS, has the potential to test, validate, and guide structure predictions of RNA:RNA complexes (*23*). While SAXS can validate the free states of the individual RNAs and the overall RNA:RNA complex, CM-SANS is necessary to validate the bound states of the individual RNAs where multiple solutions exist to satisfy the complex SAXS data. While for this simple model system that undergoes only minimal structural perturbations in the ground state, the use of the AF3 model and refinement with RNAMasonry while maintaining the kissing loop interface does result in a model from just the SAXS data that is somewhat similar to that of the SANS data; for systems with larger conformational changes, we expect the combined SAXS/SANS approach to be significantly superior to just SAXS alone.

However, like any technique, there are several limitations related to sample preparation, data acquisition, and analysis. For sample preparation, the RNA:RNA complex must remain stable over long time exposures, a challenge that could mitigated with advancements in SEC-SANS (*83*). Additionally, transient RNA:RNA interactions could be stabilized via crosslinking methods (*84*). SANS experiments are relatively sample intensive, typically requiring approximately 200–500 µL of sample at concentrations of 1-10 mg/mL, similar to sample requirements for NMR spectroscopy or X-ray crystallography.

An important consideration for SANS experiments is the contrast between the macromolecules in solution. One challenge with studying RNA:RNA complexes by SANS is that maximum contrast between the RNAs is achieved when one species is protiated and the other is perdeuterated. However, it is not possible to contrast match a perdeuterated RNA as the scattering length density is greater than that of 100% D_2_O buffer (**Fig. 3A**). To improve the signal-to-noise ratio of SANS scattering data, it would be ideal to work in 100% D_2_O contrast match buffer, where contrast between the partially deuterated and protiated species is maximized while there is no incoherent scattering from the buffer. However, it would be important to carefully determine the percent random fractional deuteration for the RNA to be contrast matched at 100% D_2_O. Additionally, unlike SAXS, which requires relatively short exposure times, SANS requires significantly longer exposure to reduce the experimental error in the scattering profiles, allowing for the averaging of more scattering data (*85*).

Nonetheless, this study demonstrates that subtle conformational differences between RNAs in isolation and in complex can be detected through SANS measurements on selectively deuterated RNA:RNA complexes. In conjunction with SAXS and *in silico* modeling, our approach allows for the reliable construction of structural models from solution scattering data. This methodology provides a general basis for the structural investigation of RNA:RNA complexes that may not be amenable to other structural techniques or can be used to validate findings from complementary methods.

## MATERIALS AND METHODS

### Construct design and template preparation

The sequences of the RNAs in this study are provided in **Table S1A**. DNA templates, which include a T7 promoter sequence, were ordered from Integrated DNA Technologies (IDT) (**Table S6**). DNA templates were C2′-methoxy-modified at the first two positions to reduce 3′-end non-templated nucleotide addition during *in vitro* transcription (IVT) (*86*). The double-stranded DNA templates for IVTs were prepared by annealing the desired DNA oligonucleotide (40 μL, 200 μM) with an oligonucleotide corresponding to the T7 promoter sequence (20 μL, 600 μM) in a boiling water bath for 3 min, followed by slow-cooling to room temperature. The duplex was subsequently diluted to 1 mL with nuclease-free water for use in IVT reactions.

### *In vitro* transcription of RNAs and purification

IVT reaction conditions were optimized prior to preparative scale transcription. RNAs were transcribed in 1× transcription buffer (40 mM Tris base, 5 mM dithiothreitol, 1 mM spermidine, and 0.01% Triton X-100, pH = 8.5), with addition of 2-10 mM rNTPs (optimized for nucleotide composition), 15-25 mM MgCl_2_, 0.8 μM annealed DNA template, 0.2 U/mL yeast inorganic pyrophosphatase (New England BioLabs), ∼15 μM T7 RNA polymerase (prepared in-house), 0.13 U/μL RNaseOUT recombinant ribonuclease inhibitor (Invitrogen), and 20% (vol/vol) dimethyl sulfoxide (DMSO). The IVT reactions were incubated at 37 °C with shaking (75 rpm) for 3-4 h. The reactions were subsequently quenched by the addition of 14.3% (vol/vol) quench buffer (7 M Urea, 0.5 M EDTA, pH = 8.0), placed in a boiling water bath for 3 min, and then snap-cooled in an ice water bath for 3 min. Glycerol (6.25% vol/vol) was added to the quenched IVT reactions prior to loading onto a preparative scale 10-14% denaturing urea polyacrylamide gel. The gels were run at a constant wattage (20 W per gel) in 0.5× TBE buffer (44.5 mM Tris pH = 8.3, 44.5 mM borate, 1 mM EDTA) for ∼24 h, or until sufficient separation was achieved. The RNA product was visualized via UV shadowing and was excised from the gel. The RNA-containing gel slices were placed in an electroelution chamber (The Gel Company) containing 0.5× TBE buffer (120 V for ∼24 hr). The RNA was collected between a Whatman 0.45 μm ME25/21 membrane filter (Cytiva) and a SnakeSkin 3.5K MWCO dialysis tubing (Thermo Scientific). Gel slices were UV shadowed to ensure complete electroelution. The eluted RNA was concentrated, washed with 2 M high purity NaCl, and desalted into nuclease-free water using Amicon Ultra-4 centrifugal filter units (MilliporeSigma). RNA purity was checked via an analytical 10-14% denaturing urea polyacrylamide gel. RNA concentration was quantified using a NanoDrop One UV-Vis spectrophotometer (Thermo Scientific). Partially- and per-deuterated RNAs were produced as described above by replacing the rNTP mixture with the desired fraction of perdeuterated rNTPs (Cambridge Isotope Laboratories).

### Native gel electrophoresis

RNA samples (10 μM, in water) were denatured by incubating in a boiling water bath for 3 min and then refolded by snap-cooling in an ice water bath for 3 min. Folding buffer was added to a final concentration of 50 mM potassium phosphate, 1 mM MgCl_2_, 50 mM NaCl, pH = 7.5 and samples were incubated at 37 °C with shaking (75 rpm) for 30 min. The pre-folded RNAs (10 μM each) were mixed together in folding buffer and incubated (37 °C, 75 rpm, 30 min) to promote complex formation. Addition of 8.3% (vol/vol) glycerol was added to the samples prior to loading. Samples were thoroughly mixed and loaded onto an 12% native polyacrylamide gel. Gels were run at 100 V at room temperature for ∼2 h, with running buffer being mixed every ∼30 min. The native gel and running buffer were prepared to a final concentration of 50 mM HEPES, 1 mM MgCl_2_, pH = 7.5. Gels were stained using a 0.0015% (w/v) Stains-All (Thermo Scientific) in a 1.5:1 formamide:water solution and destained in water. Gels were imaged using a Gel Doc XR+ system (Bio-Rad).

### Size-exclusion chromatography

Liquid chromatography was performed in 50 mM potassium phosphate buffer, 1 mM MgCl_2_, 50 mM NaCl, pH = 7.5 at room temperature on an NGC Chromatography System (Bio-Rad) equipped with a UV/Vis detector module, monitoring absorbance at 260 nm. Analytical scale size-exclusion chromatography (SEC) of RNA samples (100 μL, 0.2-0.8 mg/mL) was performed on a Superdex 75 Increase 10/300 GL column. Preparative scale SEC of RNA samples (1 mL, 1.0-3.0 mg/mL) was performed on a HiLoad 16/600 Superdex 75 pg column. Samples were either filtered using 0.22 μm Costar Spin-X centrifuge tube filters (Corning) or centrifuged (15,000 × g, 10 min) prior to loading. A flow rate of 0.5-1 mL/min was used in all chromatographic steps.

### SEC-SAXS data acquisition

SAXS was performed at BioCAT, beamline 18ID at the Advanced Photon Source (Argonne National Laboratory, Lemont, IL) in 50 mM potassium phosphate buffer, 1 mM MgCl_2_, 50 mM NaCl, pH = 7.5 at 20 °C, with in-line SEC to separate homogeneous samples of interest from aggregates and other contaminants. RNA samples (250-500 μL, 0.8-8.8 mg/mL) were loaded onto a Superdex 75 Increase 10/300 GL column (Cytiva) and run at 0.6 mL/min by an AKTA Pure FPLC (GE) with UV monitoring at 260 nm. The eluate was passed through the 1.0 mm ID quartz capillary SAXS flow cell with ∼20 μm walls. A coflowing buffer sheath was used to separate the sample from the capillary walls, reducing radiation damage (*87*). Scattering intensity was recorded using an Eiger2 XE 9M (Dectris) detector which was placed 3.678 m from the sample giving access to a *q*-range of 0.0028 Å^-1^ to 0.42 Å^-1^. 0.5 s exposures were acquired every 1 s during elution.

### SAXS data analysis and modeling

All SAXS scattering parameters are reported according to the recommended publication guidelines for SAS studies (**Table S1**) (*88*). All SAXS data was reduced and processed using BioXTAS RAW (2.2.1) (*89*). Buffer blanks, which were used for buffer subtraction, were created by averaging regions preceding and/or following the elution peak. The buffer blanks were subtracted from exposures within the elution peak to generate 1D *I*(*q*) vs *q* curves of the RNAs used for subsequent analyses. Guinier and dimensionless Kratky analyses were carried out in BioXTAS RAW (2.2.1) (*89*). The molecular mass estimates were obtained from the Porod volume (*V_p_*) using a q-cutoff of 0.15 Å^-1^ and a macromolecule density of 0.00054 kDa/Å³.(*90*) The GNOM package within ATSAS (3.0.3) was used to determine the pair-distance distribution function [*P*(*r*)] required for molecular reconstruction (*91*, *92*). DENSS was used to calculate 3D electron density reconstructions (*49*, *93*). For DENSS, 20 reconstructions were performed in slow mode using default parameters and averaged. All electron density reconstructions are contoured at 7.5σ visualizing regions with an electron density of at least: ρ̅ + 7.5σ, where ρ̅ (e/Å^3^) is the mean electron density and σ (e/Å³) is the root mean square (RMS). Structural models of the individual RNAs were predicted using RNAMasonry (0.9.14) using their sequence and predicted secondary structure [from RNAstructure (6.3)] of the RNAs (*18*, *94*). Structures were generated in an iterative manner over 100 folding steps where the models were restrained with experimental SAXS data. During the iterative folding process, FoXS was used as a measure of the goodness-of-fit (χ^2^) of the models (*51*). Rigid body modeling was performed using the SASREF package from ATSAS (3.2.1) to model the DIS-C:DIS-Gk complex, using the RNAMasonry structures as an input (*52*, *92*). Amplitude calculations of the input PDBs for SASREF was done using CRYSOL from ATSAS (3.2.1) (*95*). Theoretical scattering profiles of the complex were back-calculated using FoXS (*50*, *51*). All alignments of high-resolution atomic models were done using the “Nucleic” matrix in MatchMaker in UCSF Chimera 1.15 unless otherwise stated (*96*). All superimpositions of high resolution atomic models on solution scattering envelopes were done in UCSF Chimera 1.15 using the Fit in Map tool unless otherwise specified (*96*).

### SANS data acquisition

All SANS scattering parameters are reported according to the recommended publication guidelines for SAS studies (**Table S1**) (*88*). SANS was performed at the CG-3 Bio-SANS instrument in the High Flux Isotope Reactor facility (Oak Ridge National Laboratory, Oak Ridge, TN) (*97*, *98*). A single standard configuration of the dual detector system was set with the Panel Scan feature of the instrument with the main detector at 7 m and the wide-angle curved detector at 1.1 m and 3.2° to sample position. The instrument settings provide a *q*-range of 0.007 Å^-1^ to 1 Å^-1^. The neutron wavelength (*λ*) was 6 Å with a relative wavelength spread (Δ*λ*/*λ*) of 13.2%. SANS data were corrected for instrument background, detector sensitivity, and instrument geometry using facility data reduction software, drt-SANS (*99*). To improve data statistics and increase the number of *q* points for Guinier and *P*(*r*) analysis of the SANS data, 33 and 66 *q* values per log decade were used for the scattering vector (*q*) for samples in 65% D_2_O and 90% D_2_O buffers, respectively. Reference measurements including the direct beam, empty cell, and exactly matched buffers (obtained after buffer exchange, from either the flow-through of centrifugal ultrafiltration devices or from the SEC buffer) were measured to perform data reduction. SANS measurements were performed at room temperature with SANS samples (300 μL, 4.2-10.0 mg/mL) in 65% or 90% D_2_O (50 mM potassium phosphate buffer, 1 mM MgCl_2_, 50 mM NaCl, pH = 7.5) contained in cylindrical cells with a path length of 1 mm (Hellma).

### SANS data analysis and modeling

All further data analysis was performed using BioXTAS RAW (2.2.1) (*89*). The radii of gyration (*R_g_*) were extracted via Guinier approximation. The *P*(*r*) functions were determined using the GNOM package in ATSAS (3.0.3) (*91*, *92*). Atomic models of the RNA molecules were predicted using RNAMasonry (0.9.14) as described in the “SAXS data analysis and modeling” section except using the experimental SANS curves as folding restraints, which utilizes CRYSON from ATSAS to calculate χ^2^ (*18*, *100*). Rigid body modeling and calculation of the theoretical scattering profile of the SANS-derived complex were done as described in the “SAXS data analysis and modeling” section.

### Structure building and modeling the DIS-C:DIS-Gk kissing complex

Five models of the DIS-C:DIS-Gk kissing complex were generated using AF3 (*21*). The monomeric DIS-C and DIS-Gk SAS-derived models as described in the “SAXS/SANS data analysis and modeling” sections were aligned to the AF3 complex model 4. The apical loops of the AF3 RNAs and stems of the SAS-derived RNAs were isolated by deleting all other nucleotides. The apical loops of the AF3 RNAs were appended to the SAS-derived models using the “Adjust Bonds” tool, connecting the phosphate backbone atoms. These structural models, containing proper kissing loop (KL) stacking, were used as the input for a subsequent DIS + KL structure build in RNAMasonry (0.9.14) (*18*). In addition to the above parameters (100 folding steps, refined with the SAS data), the “freeze” command was applied to the KL residues (13-18 for DIS-C, 32-37 for DIS-Gk) of the input models, which fixed the position of those residues while allowing the other residues to be refined. To model the DIS-C:DIS-Gk complex, the individual KL models were aligned to the KLs of the AF3 complex model 4. SREFLEX was used to estimate the flexibility and optimize the top complex model against the SAXS data (*60*).

### RNA structure drawing

All RNA secondary structures were rendered using RNAcanvas (*101*).

## Data Availability

The SAXS and SANS data have been deposited at the SASBDB (*102*, *103*) and have been assigned the following accession codes: SASDUS8 (DIS-C / SAXS), SASDUT8 (DIS-Gk / SAXS), SASDUU8 (DIS-C:DIS-Gk / SAXS), SASDUV8 (^2^H-DIS-C:^1^H-DIS-Gk / SANS, 65% D_2_O), SASDUW8 (^1^H-DIS-C:^2^H-(42%)-DIS-Gk / SANS, 90% D_2_O), and SASDUX8 (^2^H(42%)-DIS-C:^1^H-DIS-Gk / SANS, 90% D_2_O).

## Acknowledgements

We are grateful to the staff at BioCAT (beamline 18ID, APS, ANL) and Bio-SANS (CG-3, HFIR, ORNL). We are also grateful to Dr. Grzegorz Chojnowski for assistance using RNAMasonry with SANS data. This research was supported by the National Science foundation (MCB-1942398, to S.C.K.), Oak Ridge Associate Universities (Ralph E. Power Junior Faculty Enhancement award, to S.C.K), National Institutes of Health Chemistry-Biology Interface training program (T32 GM132046, to A.M), and the Rackham Graduate School (Rackham Merit Fellowship, to A.M.). This research used resources of the Advanced Photon Source, a U.S. Department of Energy (DOE) Office of Science User Facility operated for the DOE Office of Science by Argonne National Laboratory under Contract No. DE-AC02-06CH11357. BioCAT was supported by grant P30 GM138395 from the National Institute of General Medical Sciences of the National Institutes of Health. This research used resources at the High Flux Isotope Reactor, a DOE Office of Science User Facility operated by the Oak Ridge National Laboratory. SANS studies were performed using the Bio-SANS instrument of the Center for Structural Molecular Biology (FWP ERKP291), a DOE Office of Biological and Environmental Research (OBER) Structural Biology Resource. The beam time was allocated to proposal numbers IPTS-30007.1 and IPTS-31445.1. The content is solely the responsibility of the authors and does not necessarily reflect the official views of the National Institute of General Medical Sciences or the National Institutes of Health.

## References

1. L. R. Ganser, M. L. Kelly, D. Herschlag, H. M. Al-Hashimi, The roles of structural dynamics in the cellular functions of RNAs. Nat Rev Mol Cell Biol 20, 474–489 (2019).

2. T. R. Cech, J. A. Steitz, The noncoding RNA revolution-trashing old rules to forge new ones. Cell 157, 77–94 (2014).

3. Y. Xue, Architecture of RNA-RNA interactions. Curr Opin Genet Dev 72, 138–144 (2022).

4. D. Wang, R. Ye, Z. Cai, Y. Xue, Emerging roles of RNA-RNA interactions in transcriptional regulation. Wiley Interdiscip Rev RNA 13, e1712 (2022).

5. S. Guil, M. Esteller, RNA-RNA interactions in gene regulation: the coding and noncoding players. Trends Biochem Sci 40, 248–56 (2015).

6. B. L. Nicholson, K. A. White, Functional long-range RNA-RNA interactions in positive-strand RNA viruses. Nat Rev Microbiol 12, 493–504 (2014).

7. I. Tinoco, C. Bustamante, How RNA folds. J Mol Biol 293, 271–281 (1999).

8. P. Brion, E. Westhof, Hierarchy and dynamics of RNA folding. Annu Rev Biophys Biomol Struct 26, 113–137 (1997).

9. B. Schneider, B. A. Sweeney, A. Bateman, J. Cerny, T. Zok, M. Szachniuk, When will RNA get its AlphaFold moment? Nucleic Acids Res 51, 9522–9532 (2023).

10. H. M. Berman, J. Westbrook, Z. Feng, G. Gilliland, T. N. Bhat, H. Weissig, I. N. Shindyalov, P. E. Bourne, The Protein Data Bank. Nucleic Acids Res 28, 235–42 (2000).

11. D. A. Jacques, J. Trewhella, Small-angle scattering for structural biology--expanding the frontier while avoiding the pitfalls. Protein Sci 19, 642–657 (2010).

12. T. M. Weiss, Small Angle Scattering: Historical Perspective and Future Outlook. Adv Exp Med Biol 1009, 1–10 (2017).

13. A. T. Tuukkanen, A. Spilotros, D. I. Svergun, Progress in small-angle scattering from biological solutions at high-brilliance synchrotrons. IUCrJ 4, 518–528 (2017).

14. J. Lipfert, S. Doniach, Small-angle X-ray scattering from RNA, proteins, and protein complexes. Annu Rev Biophys Biomol Struct 36, 307–327 (2007).

15. W. A. Cantara, E. D. Olson, K. Musier-Forsyth, Analysis of RNA structure using small-angle X-ray scattering. Methods 113, 46–55 (2017).

16. Y. Chen, L. Pollack, SAXS studies of RNA: structures, dynamics, and interactions with partners. Wiley Interdiscip Rev RNA 7, 512–526 (2016).

17. X. Fang, J. Gallego, Y.-X. Wang, Deriving RNA topological structure from SAXS. Methods Enzymol 677, 479–529 (2022).

18. G. Chojnowski, R. Zaborowski, M. Magnus, S. Mukherjee, J. M. Bujnicki, RNA 3D structure modeling by fragment assembly with small-angle X-ray scattering restraints. Bioinformatics 39, btad527 (2023).

19. M. J. Gajda, D. Martinez Zapien, E. Uchikawa, A.-C. Dock-Bregeon, Modeling the structure of RNA molecules with small-angle X-ray scattering data. PLoS One 8, e78007 (2013).

20. S. Yang, M. Parisien, F. Major, B. Roux, RNA structure determination using SAXS data. J Phys Chem B 114, 10039–10048 (2010).

21. J. Abramson, J. Adler, J. Dunger, R. Evans, T. Green, A. Pritzel, O. Ronneberger, L. Willmore, A. J. Ballard, J. Bambrick, S. W. Bodenstein, D. A. Evans, C.-C. Hung, M. O’Neill, D. Reiman, K. Tunyasuvunakool, Z. Wu, A. Žemgulytė, E. Arvaniti, C. Beattie, O. Bertolli, A. Bridgland, A. Cherepanov, M. Congreve, A. I. Cowen-Rivers, A. Cowie, M. Figurnov, F. B. Fuchs, H. Gladman, R. Jain, Y. A. Khan, C. M. R. Low, K. Perlin, A. Potapenko, P. Savy, S. Singh, A. Stecula, A. Thillaisundaram, C. Tong, S. Yakneen, E. D. Zhong, M. Zielinski, A. Žídek, V. Bapst, P. Kohli, M. Jaderberg, D. Hassabis, J. M. Jumper, Accurate structure prediction of biomolecular interactions with AlphaFold 3. Nature, doi: 10.1038/s41586-024-07487-w (2024).

22. C. Bernard, G. Postic, S. Ghannay, F. Tahi, Has AlphaFold 3 reached its success for RNAs? [Preprint] (2024). 10.1101/2024.06.13.598780.

23. N. B. Chinnam, A. Syed, G. L. Hura, M. Hammel, J. A. Tainer, S. E. Tsutakawa, Combining small angle X-ray scattering (SAXS) with protein structure predictions to characterize conformations in solution. Methods Enzymol 678, 351–376 (2023).

24. M. Sonntag, P. K. A. Jagtap, B. Simon, M.-S. Appavou, A. Geerlof, R. Stehle, F. Gabel, J. Hennig, M. Sattler, Segmental, Domain-Selective Perdeuteration and Small-Angle Neutron Scattering for Structural Analysis of Multi-Domain Proteins. Angew Chem Int Ed Engl 56, 9322–9325 (2017).

25. S.-F. Torabi, Y.-L. Chen, K. Zhang, J. Wang, S. J. DeGregorio, A. T. Vaidya, Z. Su, S. A. Pabit, W. Chiu, L. Pollack, J. A. Steitz, Structural analyses of an RNA stability element interacting with poly(A). Proc Natl Acad Sci USA 118, e2026656118 (2021).

26. T. Schamber, O. Binas, A. Schlundt, A. Wacker, H. Schwalbe, Characterization of Structure and Dynamics of the Guanidine-II Riboswitch from Escherichia coli by NMR Spectroscopy and Small-Angle X-ray Scattering (SAXS). Chembiochem 23, e202100564 (2022).

27. J. F. Pardon, D. L. Worcester, J. C. Wooley, K. Tatchell, K. E. Van Holde, B. M. Richards, Low-angle neutron scattering from chromatin subunit particles. Nucleic Acids Res 2, 2163–2176 (1975).

28. D. M. Engelman, P. B. Moore, B. P. Schoenborn, Neutron scattering measurements of separation and shape of proteins in 30S ribosomal subunit of Escherichia coli: S2-S5, S5–S8, S3-S7. Proc Natl Acad Sci U S A **72**, 3888–3892 (1975).

29. H. B. Stuhrmann, J. Haas, K. Ibel, B. Wolf, M. H. Koch, R. Parfait, R. R. Crichton, New low resolution model for 50S subunit of Escherichia coli ribosomes. Proc Natl Acad Sci U S A 73, 2379–2383 (1976).

30. F. Gabel, Small-Angle Neutron Scattering for Structural Biology of Protein-RNA Complexes. Methods Enzymol 558, 391–415 (2015).

31. J. F. Ankner, W. T. Heller, K. W. Herwig, F. Meilleur, D. A. A. Myles, Neutron scattering techniques and applications in structural biology. Curr Protoc Protein Sci **Chapter** 17, Unit17.16 (2013).

32. D. P. Hoogerheide, V. T. Forsyth, K. A. Brown, Neutron scattering for structural biology. Physics Today 73, 36–42 (2020).

33. E. Mahieu, F. Gabel, Biological small-angle neutron scattering: recent results and development. Acta Crystallogr D Struct Biol 74, 715–726 (2018).

34. J. L. Thelen, W. Leite, V. S. Urban, H. M. O’Neill, A. V. Grishaev, J. E. Curtis, S. Krueger, M. M. Castellanos, Morphological Characterization of Self-Amplifying mRNA Lipid Nanoparticles. ACS Nano 18, 1464–1476 (2024).

35. A. Hicks, P. Abraham, W. Leite, Q. Zhang, K. L. Weiss, H. O’Neill, L. Petridis, J. C. Smith, SCOMAP-XD: atomistic deuterium contrast matching for small-angle neutron scattering in biology. Acta Crystallogr D Struct Biol 79, 420–434 (2023).

36. A. E. Whitten, S. Cai, J. Trewhella, *MULCh* : modules for the analysis of small-angle neutron contrast variation data from biomolecular assemblies. J Appl Crystallogr 41, 222– 226 (2008).

37. J.-C. Paillart, M. Shehu-Xhilaga, R. Marquet, J. Mak, Dimerization of retroviral RNA genomes: an inseparable pair. Nat Rev Microbiol 2, 461–472 (2004).

38. K. Lu, X. Heng, M. F. Summers, Structural determinants and mechanism of HIV-1 genome packaging. J Mol Biol 410, 609–33 (2011).

39. N. Dubois, R. Marquet, J.-C. Paillart, S. Bernacchi, Retroviral RNA Dimerization: From Structure to Functions. Front Microbiol 9, 527 (2018).

40. J. C. Paillart, E. Skripkin, B. Ehresmann, C. Ehresmann, R. Marquet, A loop-loop “kissing” complex is the essential part of the dimer linkage of genomic HIV-1 RNA. Proc Natl Acad Sci U S A 93, 5572–7 (1996).

41. M. J. Rist, J. P. Marino, Mechanism of nucleocapsid protein catalyzed structural isomerization of the dimerization initiation site of HIV-1. Biochemistry 41, 14762–14770 (2002).

42. K. Zhang, S. C. Keane, Z. Su, R. N. Irobalieva, M. Chen, V. Van, C. A. Sciandra, J. Marchant, X. Heng, M. F. Schmid, D. A. Case, S. J. Ludtke, M. F. Summers, W. Chiu, Structure of the 30 kDa HIV-1 RNA Dimerization Signal by a Hybrid Cryo-EM, NMR, and Molecular Dynamics Approach. Structure, doi: 10.1016/j.str.2018.01.001 (2018).

43. M. D. Moore, W. Fu, O. Nikolaitchik, J. Chen, R. G. Ptak, W.-S. Hu, Dimer initiation signal of human immunodeficiency virus type 1: its role in partner selection during RNA copackaging and its effects on recombination. J Virol 81, 4002–4011 (2007).

44. L. Salmon, G. M. Giambaşu, E. N. Nikolova, K. Petzold, A. Bhattacharya, D. A. Case, H. M. Al-Hashimi, Modulating RNA Alignment Using Directional Dynamic Kinks: Application in Determining an Atomic-Resolution Ensemble for a Hairpin using NMR Residual Dipolar Couplings. J Am Chem Soc 137, 12954–12965 (2015).

45. L. A. Feigin, D. I. Svergun, Structure Analysis by Small-Angle X-Ray and Neutron Scattering (Springer US, Boston, MA, 1987; http://link.springer.com/10.1007/978-1-4757-6624-0).

46. A. G. Kikhney, D. I. Svergun, A practical guide to small angle X-ray scattering (SAXS) of flexible and intrinsically disordered proteins. FEBS Lett 589, 2570–2577 (2015).

47. D. I. Svergun, Determination of the regularization parameter in indirect-transform methods using perceptual criteria. J Appl Crystallogr 25, 495–503 (1992).

48. D. I. Svergun, M. H. J. Koch, Small-angle scattering studies of biological macromolecules in solution. Rep. Prog. Phys. 66, 1735–1782 (2003).

49. T. D. Grant, Ab initio electron density determination directly from solution scattering data. Nat Methods 15, 191–193 (2018).

50. D. Schneidman-Duhovny, M. Hammel, J. A. Tainer, A. Sali, Accurate SAXS profile computation and its assessment by contrast variation experiments. Biophys J 105, 962–974 (2013).

51. D. Schneidman-Duhovny, M. Hammel, J. A. Tainer, A. Sali, FoXS, FoXSDock and MultiFoXS: Single-state and multi-state structural modeling of proteins and their complexes based on SAXS profiles. Nucleic Acids Res 44, W424–429 (2016).

52. M. V. Petoukhov, D. I. Svergun, Global rigid body modeling of macromolecular complexes against small-angle scattering data. Biophys J 89, 1237–1250 (2005).

53. N. B. Leontis, E. Westhof, Geometric nomenclature and classification of RNA base pairs. RNA 7, 499–512 (2001).

54. C. M. Jeffries, Z. Pietras, D. I. Svergun, The basics of small-angle neutron scattering (SANS for new users of structural biology). EPJ Web Conf. 236, 03001 (2020).

55. K. Lu, Y. Miyazaki, M. F. Summers, Isotope labeling strategies for NMR studies of RNA. J Biomol NMR 46, 113–25 (2010).

56. H. Zhang, S. C. Keane, Advances that facilitate the study of large RNA structure and dynamics by nuclear magnetic resonance spectroscopy. Wiley Interdiscip Rev RNA, e1541 (2019).

57. A. Kotar, H. N. Foley, K. M. Baughman, S. C. Keane, Advanced approaches for elucidating structures of large RNAs using NMR spectroscopy and complementary methods. Methods, doi: 10.1016/j.ymeth.2020.01.009 (2020).

58. K. Wood, M. Plazanet, F. Gabel, B. Kessler, D. Oesterhelt, G. Zaccai, M. Weik, Dynamics of hydration water in deuterated purple membranes explored by neutron scattering. Eur Biophys J 37, 619–626 (2008).

59. R. R. Crichton, D. M. Engleman, J. Haas, M. H. Koch, P. B. Moore, R. Parfait, H. B. Stuhrmann, Contrast variation study of specifically deuterated Escherichia coli ribosomal subunits. Proc Natl Acad Sci U S A 74, 5547–5550 (1977).

60. A. Panjkovich, D. I. Svergun, Deciphering conformational transitions of proteins by small angle X-ray scattering and normal mode analysis. Phys Chem Chem Phys 18, 5707–5719 (2016).

61. X. Sun, Q. Zhang, H. M. Al-Hashimi, Resolving fast and slow motions in the internal loop containing stem-loop 1 of HIV-1 that are modulated by Mg2+ binding: role in the kissing– duplex structural transition. Nucleic Acids Research 35, 1698–1713 (2007).

62. E. A. Dethoff, K. Petzold, J. Chugh, A. Casiano-Negroni, H. M. Al-Hashimi, Visualizing transient low-populated structures of RNA. Nature 491, 724–8 (2012).

63. M.-R. Mihailescu, J. P. Marino, A proton-coupled dynamic conformational switch in the HIV-1 dimerization initiation site kissing complex. Proc. Natl. Acad. Sci. U.S.A. 101, 1189–1194 (2004).

64. F. Y. F. Tieng, M.-R. Abdullah-Zawawi, N. A. A. Md Shahri, Z.-A. Mohamed-Hussein, L.-H. Lee, N.-S. A. Mutalib, A Hitchhiker’s guide to RNA-RNA structure and interaction prediction tools. Brief Bioinform 25, bbad421 (2023).

65. I. K. Beckmann, M. Waldl, S. Will, I. L. Hofacker, 3D feasibility of 2D RNA-RNA interaction paths by stepwise folding simulations. RNA 30, 113–123 (2024).

66. S. Singh, S. Shyamal, A. C. Panda, Detecting RNA-RNA interactome. Wiley Interdiscip Rev RNA 13, e1715 (2022).

67. T. Mrozowich, S. M. Park, M. Waldl, A. Henrickson, S. Tersteeg, C. R. Nelson, A. De Klerk, B. Demeler, I. L. Hofacker, M. T. Wolfinger, T. R. Patel, Investigating RNA-RNA interactions through computational and biophysical analysis. Nucleic Acids Res 51, 4588– 4601 (2023).

68. S. Krueger, Small-angle neutron scattering contrast variation studies of biological complexes: Challenges and triumphs. Curr Opin Struct Biol 74, 102375 (2022).

69. J. C. Grigg, Y. Chen, F. J. Grundy, T. M. Henkin, L. Pollack, A. Ke, T box RNA decodes both the information content and geometry of tRNA to affect gene expression. Proc Natl Acad Sci U S A 110, 7240–7245 (2013).

70. L. Baronti, H. Karlsson, M. Marušič, K. Petzold, A guide to large-scale RNA sample preparation. Anal Bioanal Chem 410, 3239–3252 (2018).

71. J. San Emeterio, S. A. Pabit, L. Pollack, Contrast variation SAXS: Sample preparation protocols, experimental procedures, and data analysis. Methods Enzymol 677, 41–83 (2022).

72. A. Gupta, J. L. Andresen, R. S. Manan, R. Langer, Nucleic acid delivery for therapeutic applications. Adv Drug Deliv Rev 178, 113834 (2021).

73. J. S. Smith, E. P. Nikonowicz, Phosphorothioate Substitution Can Substantially Alter RNA Conformation. Biochemistry 39, 5642–5652 (2000).

74. A. Carlesso, J. Hörberg, G. Deganutti, A. Reymer, P. Matsson, Structural dynamics of therapeutic nucleic acids with phosphorothioate backbone modifications. NAR Genomics and Bioinformatics 6, lqae058 (2024).

75. D. Liu, R. Xu, D. Cowburn, Segmental isotopic labeling of proteins for nuclear magnetic resonance. Methods Enzymol 462, 151–175 (2009).

76. M. Muona, A. S. Aranko, V. Raulinaitis, H. Iwaï, Segmental isotopic labeling of multi-domain and fusion proteins by protein trans-splicing in vivo and in vitro. Nat Protoc 5, 574–587 (2010).

77. J. Xue, D. S. Burz, A. Shekhtman, Segmental labeling to study multidomain proteins. Adv Exp Med Biol 992, 17–33 (2012).

78. J. Li, Y. Zhang, O. Soubias, D. Khago, F.-A. Chao, Y. Li, K. Shaw, R. A. Byrd, Optimization of sortase A ligation for flexible engineering of complex protein systems. J Biol Chem 295, 2664–2675 (2020).

79. J. Xu, J. Lapham, D. M. Crothers, Determining RNA solution structure by segmental isotopic labeling and NMR: application to Caenorhabditis elegans spliced leader RNA 1. Proc Natl Acad Sci U S A 93, 44–8 (1996).

80. O. Duss, C. Maris, C. von Schroetter, F. H. Allain, A fast, efficient and sequence-independent method for flexible multiple segmental isotope labeling of RNA using ribozyme and RNase H cleavage. Nucleic Acids Res 38, e188 (2010).

81. O. Duss, P. J. Lukavsky, F. H.-T. Allain, Isotope labeling and segmental labeling of larger RNAs for NMR structural studies. Adv Exp Med Biol 992, 121–144 (2012).

82. R. Haslecker, V. Pham, D. Glänzer, C. Kreutz, T. Dayie, V. D’Souza, Extending the toolbox for RNA biology with SegModTeX: A polymerase-driven method for site-specific and segmental labeling of RNA. [Preprint] (2023). 10.21203/rs.3.rs-2782805/v1.

83. N. T. Johansen, M. C. Pedersen, L. Porcar, A. Martel, L. Arleth, Introducing SEC–SANS for studies of complex self-organized biological systems. Acta Crystallogr D Struct Biol 74, 1178–1191 (2018).

84. M. E. Harris, E. L. Christian, RNA crosslinking methods. Methods Enzymol 468, 127–146 (2009).

85. M. C. Pedersen, S. L. Hansen, B. Markussen, L. Arleth, K. Mortensen, Quantification of the information in small-angle scattering data. J Appl Crystallogr 47, 2000–2010 (2014).

86. C. Kao, M. Zheng, S. Rudisser, A simple and efficient method to reduce nontemplated nucleotide addition at the 3 terminus of RNAs transcribed by T7 RNA polymerase. RNA 5, 1268–72 (1999).

87. N. Kirby, N. Cowieson, A. M. Hawley, S. T. Mudie, D. J. McGillivray, M. Kusel, V. Samardzic-Boban, T. M. Ryan, Improved radiation dose efficiency in solution SAXS using a sheath flow sample environment. Acta Crystallogr D Struct Biol 72, 1254–1266 (2016).

88. J. Trewhella, A. P. Duff, D. Durand, F. Gabel, J. M. Guss, W. A. Hendrickson, G. L. Hura, D. A. Jacques, N. M. Kirby, A. H. Kwan, J. Pérez, L. Pollack, T. M. Ryan, A. Sali, D. Schneidman-Duhovny, T. Schwede, D. I. Svergun, M. Sugiyama, J. A. Tainer, P. Vachette, J. Westbrook, A. E. Whitten, 2017 publication guidelines for structural modelling of small-angle scattering data from biomolecules in solution: an update. Acta Crystallogr D Struct Biol 73, 710–728 (2017).

89. J. B. Hopkins, BioXTAS RAW 2: new developments for a free open-source program for small-angle scattering data reduction and analysis. J Appl Crystallogr 57, 194–208 (2024).

90. V. Piiadov, E. Ares De Araújo, M. Oliveira Neto, A. F. Craievich, I. Polikarpov, SAXSMoW 2.0: Online calculator of the molecular weight of proteins in dilute solution from experimental SAXS data measured on a relative scale. Protein Science 28, 454–463 (2019).

91. A. V. Semenyuk, D. I. Svergun, GNOM – a program package for small-angle scattering data processing. J Appl Crystallogr 24, 537–540 (1991).

92. K. Manalastas-Cantos, P. V. Konarev, N. R. Hajizadeh, A. G. Kikhney, M. V. Petoukhov, D. S. Molodenskiy, A. Panjkovich, H. D. T. Mertens, A. Gruzinov, C. Borges, C. M. Jeffries, D. I. Svergun, D. Franke, ATSAS 3.0: expanded functionality and new tools for small-angle scattering data analysis. J Appl Crystallogr 54, 343–355 (2021).

93. D. Franke, D. I. Svergun, DAMMIF, a program for rapid ab-initio shape determination in small-angle scattering. J Appl Crystallogr 42, 342–346 (2009).

94. S. Bellaousov, J. S. Reuter, M. G. Seetin, D. H. Mathews, RNAstructure: web servers for RNA secondary structure prediction and analysis. Nucleic Acids Research 41, W471– W474 (2013).

95. D. Svergun, C. Barberato, M. H. J. Koch, *CRYSOL* – a Program to Evaluate X-ray Solution Scattering of Biological Macromolecules from Atomic Coordinates. J Appl Crystallogr 28, 768–773 (1995).

96. E. F. Pettersen, T. D. Goddard, C. C. Huang, G. S. Couch, D. M. Greenblatt, E. C. Meng, T. E. Ferrin, UCSF Chimera--a visualization system for exploratory research and analysis. J Comput Chem 25, 1605–1612 (2004).

97. W. T. Heller, V. S. Urban, G. W. Lynn, K. L. Weiss, H. M. O’Neill, S. V. Pingali, S. Qian, K. C. Littrell, Y. B. Melnichenko, M. V. Buchanan, D. L. Selby, G. D. Wignall, P. D. Butler, D. A. Myles, The Bio-SANS instrument at the High Flux Isotope Reactor of Oak Ridge National Laboratory. J Appl Crystallogr 47, 1238–1246 (2014).

98. W. T. Heller, Small-angle neutron scattering and contrast variation: a powerful combination for studying biological structures. Acta Crystallogr D Biol Crystallogr 66, 1213–1217 (2010).

99. W. T. Heller, J. Hetrick, J. Bilheux, J. M. B. Calvo, W.-R. Chen, L. DeBeer-Schmitt, C. Do, M. Doucet, M. R. Fitzsimmons, W. F. Godoy, G. E. Granroth, S. Hahn, L. He, F. Islam, J. Lin, K. C. Littrell, M. McDonnell, J. McGaha, P. F. Peterson, S. V. Pingali, S. Qian, A. T. Savici, Y. Shang, C. B. Stanley, V. S. Urban, R. E. Whitfield, C. Zhang, W. Zhou, J. J. Billings, M. J. Cuneo, R. M. F. Leal, T. Wang, B. Wu, drtsans: The data reduction toolkit for small-angle neutron scattering at Oak Ridge National Laboratory. SoftwareX 19, 101101 (2022).

100. D. I. Svergun, S. Richard, M. H. Koch, Z. Sayers, S. Kuprin, G. Zaccai, Protein hydration in solution: experimental observation by x-ray and neutron scattering. Proc Natl Acad Sci U S A 95, 2267–2272 (1998).

101. P. Z. Johnson, A. E. Simon, RNAcanvas: interactive drawing and exploration of nucleic acid structures. Nucleic Acids Research 51, W501–W508 (2023).

102. E. Valentini, A. G. Kikhney, G. Previtali, C. M. Jeffries, D. I. Svergun, SASBDB, a repository for biological small-angle scattering data. Nucleic Acids Res 43, D357–363 (2015).

103. A. G. Kikhney, C. R. Borges, D. S. Molodenskiy, C. M. Jeffries, D. I. Svergun, SASBDB: Towards an automatically curated and validated repository for biological scattering data. Protein Sci 29, 66–75 (2020).

104. D. Franke, C. M. Jeffries, D. I. Svergun, Correlation Map, a goodness-of-fit test for one-dimensional X-ray scattering spectra. Nat Methods 12, 419–422 (2015).

105. D. Svergun, C. Barberato, M. H. Koch, CRYSOL - A program to evaluate X-ray solution scattering of biological macromolecules from atomic coordinates. Journal of Applied Crystallography 28, 768–773 (1995).

