## Supplementary Data for "Selective deuteration of an RNA:RNA complex for structural analysis using small-angle scattering"

**Table S1. SAXS and SANS parameters for DIS RNA samples according to publication guidelines.**(*88*)
**(a) Sample details.**

| Sample<br>**Visible:***Matched*<br>(% $D_2O$ buffer) | DIS-C<br>SAXS | DIS-Gk<br>SAXS | DIS-C:<br>DIS-Gk<br>SAXS | **$^{2}$H-DIS-C:**<br>*$^{1}$H-DIS-Gk*<br>SANS (65%) | **$^{1}$H-DIS-C:**<br>*$^{2}$H(42%)-DIS-Gk*<br>SANS (90%) | *$^{2}$H(42%)-DIS-C*:<br>**$^{1}$H-DIS-Gk**<br>SANS (90%) |
|---|---|---|---|---|---|---|
| Organism | Human immunodeficiency virus type 1 | | | | | |
| Source | *In vitro* transcribed RNA | | | | | |
| Sequence (5′ to 3′) | GGGCUUGC<br>UGAACCCCC<br>CACGGCAA<br>GACC | GGGUCGCG<br>CUGGCAGA<br>UCUGGGCU<br>UGCUGAAG<br>GGGGGACG<br>GCAAGACC<br>UCUGCCAG<br>CGCGACCC | 1:1 DIS-C:DIS-Gk heterodimer | | | |
| Extinction coefficient ($M^{-1}$ $cm^{-1}$) | 266000 | 600800 | 866800 | | | |
| $\bar{v}$ from chemical composition (mL/g) | 0.5696 | 0.5687 | 0.569 | | | |
| Particle contrast from sequence and solvent constituents, $\Delta\overline{\rho}$ ($\rho_{RNA} - \rho_{solvent}$) (Å$^{-1}$) | - | - | - | 3.38 | -1.73 | -1.73 |
| *M* from chemical composition (kDa) | 9.34 | 20.84 | 30.18 | | | |
| SEC-SAXS column | Superdex 75 10/300 Increase | | | - | | |
| Sample concentration (mg $mL^{-1}$) | 8.8 | 0.8 | 3.4 | 4.2 | 10.0 | 4.5 |
| Volume (µL) | 250 | 500 | 250 | 300 | 300 | 300 |
| Flow rate (mL/min) | 0.6 | 0.6 | 0.6 | - | | |
| Solvent | 50 mM potassium phosphate buffer, 1 mM $MgCl_2$, 50 mM NaCl, pH = 7.5 | | | 50 mM potassium phosphate buffer, 1 mM $MgCl_2$, 50 mM NaCl, pH = 7.5 (65% $D_2O$) | 50 mM potassium phosphate buffer, 1 mM $MgCl_2$, 50 mM NaCl, pH = 7.5 (90% $D_2O$) | 50 mM potassium phosphate buffer, 1 mM $MgCl_2$, 50 mM NaCl, pH = 7.5 (90% $D_2O$) |

**(b) SAXS/SANS data collection parameters.**

| | SAXS | SANS |
|---|---|---|
| Instrument/data processing | BioCAT (beamline 18ID, APS) with a Dectris Eiger2 XE 9M detector | Bio-SANS (CG-3, HFIR) with a 2D linear position sensitive detector |
| Wavelength (Å) | 1.033 | 6.0 |
| Beam size (µm) | 30 × 150 (focused on detector) | 76000 (focused on detector) |
| Camera length (m) | 3.678 | 7 (main detector), 1.1 (sample-to-detector distance and 3.2° rotation, wide angle detector) |
| $q$ measurement range (Å$^{-1}$) | 0.0028 – 0.42 | 0.007 – 1.0 |
| Absolute scaling method | Glassy Carbon, NIST SRM 3600 | Porous Silica |
| Normalization | To transmitted intensity by beam-stop counter | To incident intensity by monitor beam counter |
| Monitoring for radiation damage | Automated frame-by-frame comparison of relevant regions using CORMAP implemented in BioXTAS RAW(*89*, *104*) | No radiation damage |
| Exposure time | 0.5 s exposure time with a 1 s total exposure period (0.5 s on, 0.5 s off) of entire SEC elution | 37800 – 68400 s (sample, buffer, empty cell) |
| Sample configuration | SEC–SAXS with sheath-flow cell, effective path length 0.542 mm(*87*) | Batch mode SANS |
| Sample temperature (°C) | 20 | |

**(c) Software employed for SAXS/SANS data reduction, analysis, and interpretation.**

| | **SAXS** | **SANS** |
|---|---|---|
| Data reduction | *I*(*q*) vs *q* and solvent subtraction using BioXTAS RAW 2.2.1(*89*) | *I*(*q*) vs *q* using drt-SANS, solvent subtraction using BioXTAS RAW 2.2.1(*89*, *99*) |
| Extinction coefficient estimate | Quest Calculate™ RNA Concentration Calculator *via* web server (https://www.aatbio.com/tools/calculate-RNA-concentration) | |
| Basic analyses, Guinier, $P(r)$, $V_p$ | BioXTAS RAW 2.2.1 and GNOM from ATSAS 3.0.3(*89*, *92*) | |
| Electron density modeling | DENSS(*49*) | - |
| Atomic structure modeling | RNAMasonry (0.9.14)(*18*)<br>FoXS *via* web server (https://modbase.compbio.ucsf.edu/foxs/)(*51*)<br>CRYSON from ATSAS 3.2.1(*92*)<br>SASREF from ATSAS 3.2.1(*52*)<br>CRYSOL from ATSAS 3.2.1(*105*)<br>SREFLEX from ATSAS 3.2.1(*60*) | |
| Three-dimensional graphic model representations | UCSF Chimera 1.15(*96*) | |

**(d) Structural parameters**

| Sample **Visible:***Matched* (% $D_2O$ buffer) | DIS-C SAXS | DIS-Gk SAXS | DIS-C:DIS-Gk SAXS | **$^2$H-DIS-C:** *$^1$H-DIS-Gk* SANS (65%) | **$^1$H-DIS-C:** *$^2$H(42%)-DIS-Gk* SANS (90%) | *$^2$H(42%)-DIS-C*: **$^1$H-DIS-Gk** SANS (90%) |
|---|---|---|---|---|---|---|
| **Guinier analysis** | | | | | | |
| $I(0)$ ($cm^{-1}$) | 0.03 ± 2.53×10$^{-5}$ | 0.02 ± 3.71×10$^{-5}$ | 0.03 ± 4.36×10$^{-5}$ | 1.8 ± 0.1 (×10$^{-3}$) | 1.7 ± 0.2 (×10$^{-3}$) | 3.4 ± 0.3 (×10$^{-3}$) |
| $R_g$ (Å) | 14.65 ± 0.04 | 26.4 ± 0.1 | 35.3 ± 0.1 | 13.3 ± 1.8 | 14.1 ± 3.5 | 26.7 ± 3.3 |
| $q_{min}$ (Å$^{-1}$) | 0.00279 | 0.00279 | 0.00279 | 0.040 | 0.008 | 0.008 |
| $qR_g$ max | 1.0070 | 1.1173 | 1.0030 | 1.32 | 1.26 | 1.32 |
| Coefficient of correlation, $R^2$ | 0.9813 | 0.9563 | 0.9904 | * | * | * |
| $M$ from $V_p$ (kDa) | 7.9 | 21.8 | 30.8 | 9.8 | 10.6 | 19.9 |
| $V_p$ (Å$^3$) | 4.28×10$^4$ | 8.74×10$^4$ | 1.16×10$^5$ | 4.85×10$^4$ | 5.12×10$^4$ | 8.13×10$^4$ |
| ***P*(*r*) analysis** | | | | | | |
| $I(0)$ ($cm^{-1}$) | 0.03 ± 1.89×10$^{-5}$ | 0.02 ± 3.8×10$^{-5}$ | 0.03 ± 4.04×10$^{-5}$ | 1.9 ± 0.1 (×10$^{-3}$) | 1.5 ± 0.1 (×10$^{-3}$) | 3.0 ± 0.2 (×10$^{-3}$) |
| $R_g$ (Å) | 15.12 ± 0.02 | 27.5 ± 0.1 | 36.5 ± 0.1 | 16.3 ± 1.3 | 16.8 ± 1.5 | 26.4 ± 1.3 |
| $D_{max}$ (Å) | 46 | 105 | 135 | 53 | 46 | 97 |
| $q$ range (Å$^{-1}$) | 0.0028 – 0.3002 | 0.0028 – 0.3002 | 0.0028 – 0.3002 | 0.0374 – 0.3736 | 0.0193 – 0.2485 | 0.007 – 0.1617 |
| $\chi^2$ | 1.21 | 1.07 | 0.91 | 0.73 | 1.04 | 0.78 |

*Due the low signal-to-noise ratio in the low-q regime, Guinier fits for the SANS data did not provide reliable coefficient of correlations.

### (e) Shape model-fitting results

| Sample | DIS-C SAXS | DIS-Gk SAXS | DIS-C:DIS-Gk SAXS |
|---|---|---|---|
| **DENSS (default parameters, 20 calculations)** | | | |
| $q$-range for fitting (Å$^{-1}$) | 0.0028 – 0.3002 | 0.0028 – 0.3002 | 0.0028 – 0.3002 |
| Symmetry, anisotropy assumption | P1, none | | |
| Ambiguity score (AMBIMETER) | 1.699 | 2.430 | 2.161 |
| $\chi^2$ range | 0.00003 - 0.01455 | 0.00024 - 0.02013 | 0.0072 - 0.37433 |
| Model resolution (Å) | 19.0 | 23.6 | 30.1 |

#### (f) Atomistic modeling for DIS-C

| Model | DIS-C SAXS | DIS-C SAXS + KL | DIS-C SANS (65% $D_2O$) | DIS-C SANS + KL (65% $D_2O$) | DIS-C SANS (90% $D_2O$) | DIS-C SANS + KL (90% $D_2O$) |
|---|---|---|---|---|---|---|
| **RNAMasonry** | | | | | | |
| Input sequence/structure | DIS-C sequence in **(a)** | Structure S2 (provided in SI) | DIS-C sequence in **(a)** | Structure S5 (provided in SI) | DIS-C sequence in **(a)** | Structure S8 (provided in SI) |
| SAS curve used to restrain folding | SASDUS8 | SASDUS8 | SASDUV8 | SASDUV8 | SASDUW8 | SASDUW8 |
| *q*-range for folding ($Å^{-1}$) | 0.0028 – 0.4157 | | 0.0099 - 0.9254 | | 0.007 - 0.9254 | |
| Software used to calculate $\chi^2$ during folding | FoXS(*50*) | | CRYSON from ATSAS 3.2.1(*92*) | | | |
| Residues frozen | | 13:18 | | 13:18 | | 13:18 |
| CRYSON options | - | - | -D2O 0.65 -per 1.0 | -D2O 0.65 -per 1.0 | -D2O 0.90 -per 0.0 | -D2O 0.90 -per 0.0 |
| Number of folding steps | 100 | | | | | |
| $\chi^2$ of final model | 1.23 | 2.95 | 1.34 | 1.22 | 0.79 | 0.79 |

### (g) Atomistic modeling for DIS-Gk

| **Model** | **DIS-Gk SAXS** | **DIS-Gk SAXS + KL** | **DIS-Gk SANS (90% $D_2O$)** | **DIS-Gk SANS + KL (90% $D_2O$)** |
|---|---|---|---|---|
| **RNAMasonry** | | | | |
| Input sequence/structure | DIS-Gk sequence in **(a)** | Structure S11 (provided in SI) | DIS-Gk sequence in **(a)** | Structure S14 (provided in SI) |
| SAS curve used to restrain folding | SASDUT8 | SASDUT8 | SASDUX8 | SASDUX8 |
| *q*-range for folding ($Å^{-1}$) | 0.0028 – 0.4157 | | 0.007 - 0.9254 | |
| Software used to calculate $\chi^2$ during folding | FoXS(*50*) | | CRYSON from ATSAS 3.2.1(*92*) | |
| Residues frozen | - | 32:37 | - | 32:37 |
| CRYSON options | - | - | -D2O 0.90 -per 0.0 | -D2O 0.65 -per 0.0 |
| Number of folding steps | 100 | | | |
| $\chi^2$ of final model | 1.13 | 0.99 | 0.74 | 0.77 |

**(h) SASBDB IDs for data and models**(*102*, *103*)

| Sample<br>**Visible:*****Matched***<br>(% $D_2O$ buffer) | DIS-C<br>SAXS | DIS-Gk<br>SAXS | DIS-C:DIS-Gk<br>SAXS | $^{2}$**H-DIS-C:**<br>$^{1}$***H-DIS-Gk***<br>SANS (65%) | $^{1}$**H-DIS-C:**<br>$^{2}$***H(42%)-DIS-Gk***<br>SANS (90%) | $^{2}$***H(42%)-DIS-C*****:**<br>$^{1}$**H-DIS-Gk**<br>SANS (90%) |
|---|---|---|---|---|---|---|
| SASBDB ID | SASDUS8 | SASDUT8 | SASDUU8 | SASDUV8 | SASDUW8 | SASDUX8 |

**Table S2. DNA oligonucleotides used in this work.**

| Oligo name | Sequence (5′ to 3′)[a,b] |
|---|---|
| T7 promoter | <u>TAATACGACTCACTATA</u> |
| DIS-C | mGmGTCTTGCCGTGGGGGGTTCAGCAAGCCC<u>TATAGTGAGTCGTATTA</u> |
| DIS-Gk | mGmGGTCGCGCTGGCAGAGGTCTTGCCGTCCCCCCTTCAGCAAGCCCAGATCTGCCAGCGCGACCC<u>TATAGTGAGTCGTATTA</u> |

[a] m denotes 2′-O-Methyl modification of the nucleotide.

[b] Underlined nucleotides indicate the T7 promoter sequence.

**Table S3. Comparison of the DIS-C RNA models**

| **Model** | **DIS-C SAXS** | **DIS-C SAXS + KL** | **DIS-C SANS (65% $D_2O$)** | **DIS-C SANS + KL (65% $D_2O$)** | **DIS-C SANS (90% $D_2O$)** | **DIS-C SANS + KL (90% $D_2O$)** | ***q*-range used for fitting ($Å^{-1}$)** |
|---|---|---|---|---|---|---|---|
| FoXS theoretical $R_g$ (Å) | 14.65 | 14.79 | 14.19 | 14.57 | 14.90 | 15.02 | - |
| FoXS goodness-of-fit to DIS-C SAXS data ($\chi^2$) | 1.23 | 2.95 | 2.05 | 4.33 | 3.57 | 2.53 | 0.0028 - 0.4157 |
| CRYSON goodness-of-fit to DIS-C SANS (65%) data ($\chi^2$) | 1.34 | 1.41 | 1.34 | 1.22 | 1.40 | 1.38 | 0.0099 - 0.9254 |
| CRYSON goodness-of-fit to DIS-C SANS (90%) data ($\chi^2$) | 0.79 | 0.78 | 0.79 | 0.80 | 0.79 | 0.79 | 0.007 - 0.9254 |

**Table S4. Comparison of the DIS-Gk RNA models**

| Model | DIS-Gk SAXS | DIS-Gk SAXS + KL | DIS-Gk SANS (90% $D_2O$) | DIS-Gk SANS + KL (90% $D_2O$) | *q*-range used for fitting (Å$^{-1}$) |
|---|---|---|---|---|---|
| FoXS theoretical $R_g$ (Å) | 26.32 | 26.46 | 25.54 | 26.99 | - |
| FoXS goodness-of-fit to DIS-Gk SAXS data ($\chi^2$) | 1.13 | 0.99 | 1.88 | 2.57 | 0.0028 - 0.4157 |
| CRYSON goodness-of-fit to DIS-Gk SANS (90%) data ($\chi^2$) | 0.75 | 0.74 | 0.74 | 0.77 | 0.007 - 0.9254 |

**Table S5. Comparison of AlphaFold 3 (AF3) DIS-C:DIS-Gk kissing complex models**

| **Model** | **1** | **2** | **3** | **4** | **5** | ***q*-range used for fitting (Å$^{-1}$)** |
|---|---|---|---|---|---|---|
| FoXS theoretical $R_g$ (Å) | 35.02 | 35.11 | 35.66 | 34.34 | 34.33 | - |
| FoXS goodness-of-fit to DIS-C:DIS-Gk SAXS data ($\chi^2$) | 6.22 | 3.72 | 3.77 | 3.66 | 5.76 | 0.0028 - 0.4157 |

**Table S6. Comparison of SAS-derived DIS-C:DIS-Gk kissing complex models**

| **Model** | **DIS-C + KL (SAXS):DIS-Gk + KL (SAXS)** | **DIS-C + KL (SAXS):DIS-Gk + KL (SANS, 90% $D_2O$)** | **DIS-C + KL (SANS, 65% $D_2O$):DIS-Gk + KL (SAXS)** | **DIS-C + KL (SANS, 65% $D_2O$):DIS-Gk + KL (SANS, 90% $D_2O$)** | **DIS-C + KL (SANS, 90% $D_2O$):DIS-Gk + KL (SAXS)** | **DIS-C + KL (SANS, 90% $D_2O$):DIS-Gk + KL (SANS, 90% $D_2O$)** | ***q*-range used for fitting ($Å^{-1}$)** |
|---|---|---|---|---|---|---|---|
| FoXS theoretical $R_g$ (Å) | 35.08 | 35.58 | 35.50 | 35.88 | 35.24 | 35.74 | - |
| FoXS goodness-of-fit to DIS-C:DIS-Gk SAXS data ($\chi^2$) | 1.30 | 2.98 | 1.11 | 1.98 | 1.20 | 2.99 | 0.0028 - 0.4157 |

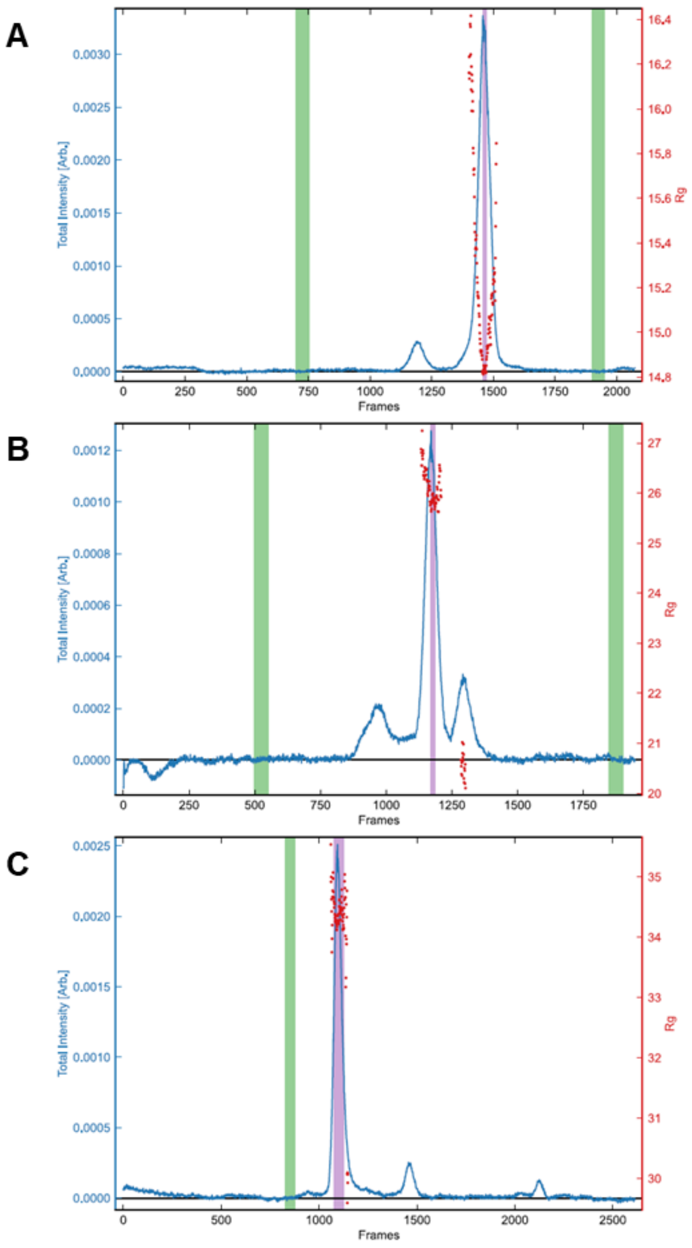


**Figure S1. SEC-SAXS profiles of DIS-C, DIS-Gk, and DIS-C:DIS-Gk.** Series intensity (blue, left axis) vs. frame and $R_g$ vs. frame (red, right axis). Green shaded regions are buffer regions, purple shaded regions are sample regions for **(A)** DIS-C, **(B)** DIS-Gk, and **(C)** DIS-C:DIS-Gk.

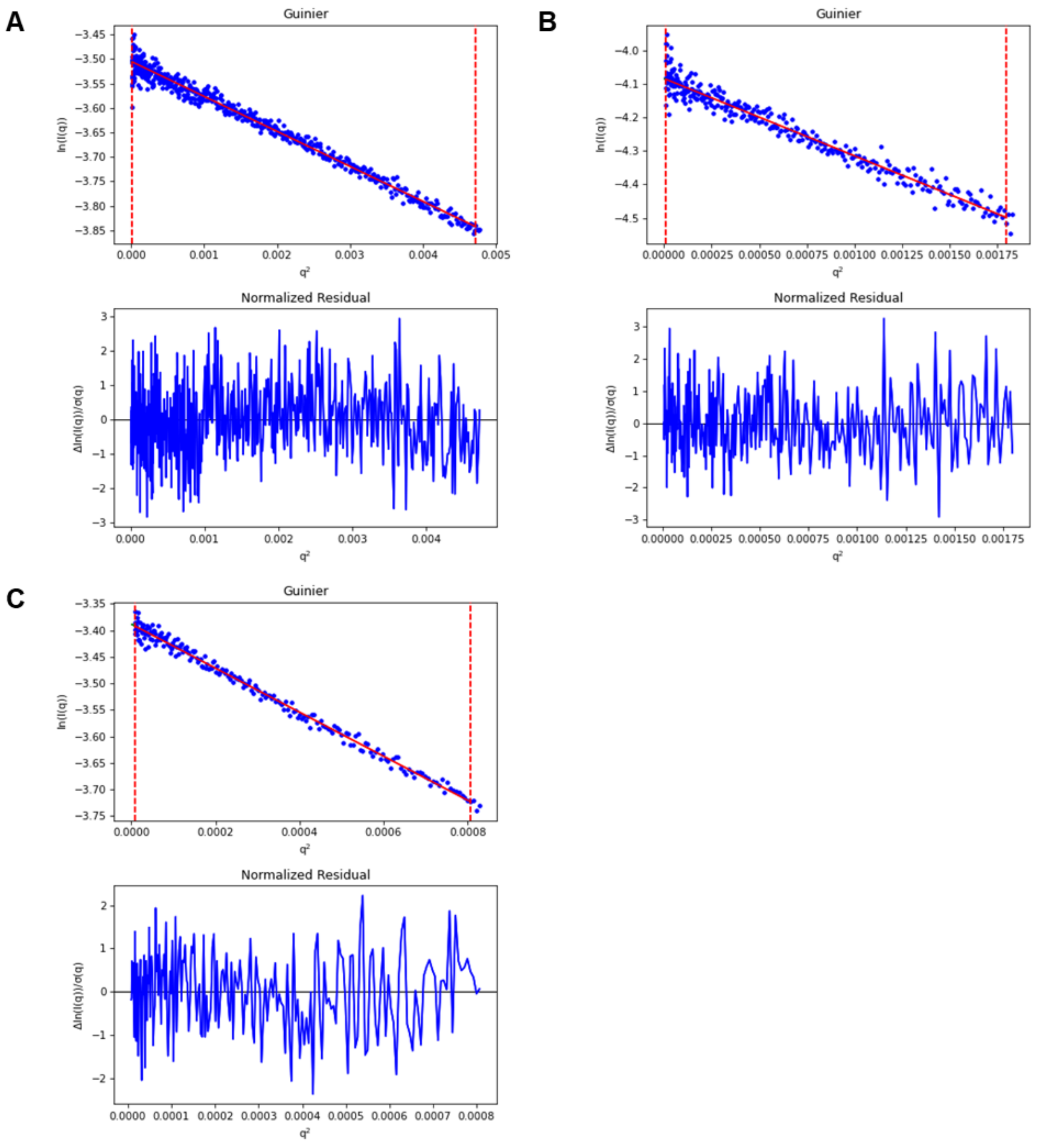


**Figure S2. SAXS Guinier analysis of DIS-C, DIS-Gk, and DIS-C:DIS-Gk.** Guinier fits (red) to experimental data (blue) for **(A)** DIS-C, **(B)** DIS-Gk, and **(C)** DIS-C:DIS-Gk with corresponding residuals.

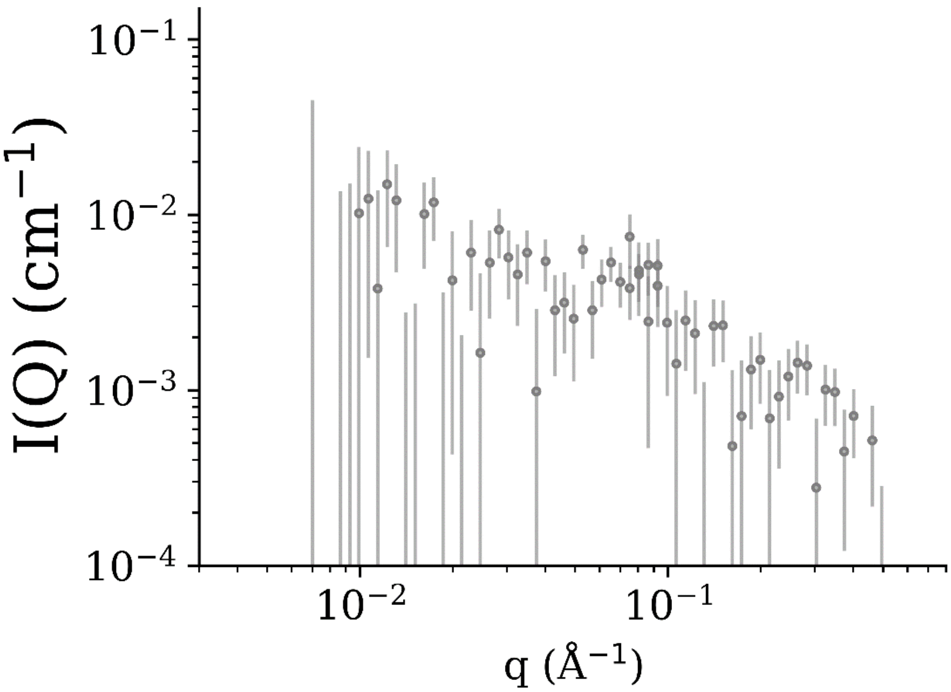


**Figure S3. $^{2}$H(42%)-DIS-C RNA is contrast matched at 90% $D_2O$ buffer.** Experimental SANS of $^{2}$H(42%)-DIS-C in 90% $D_2O$ buffer (match buffer).

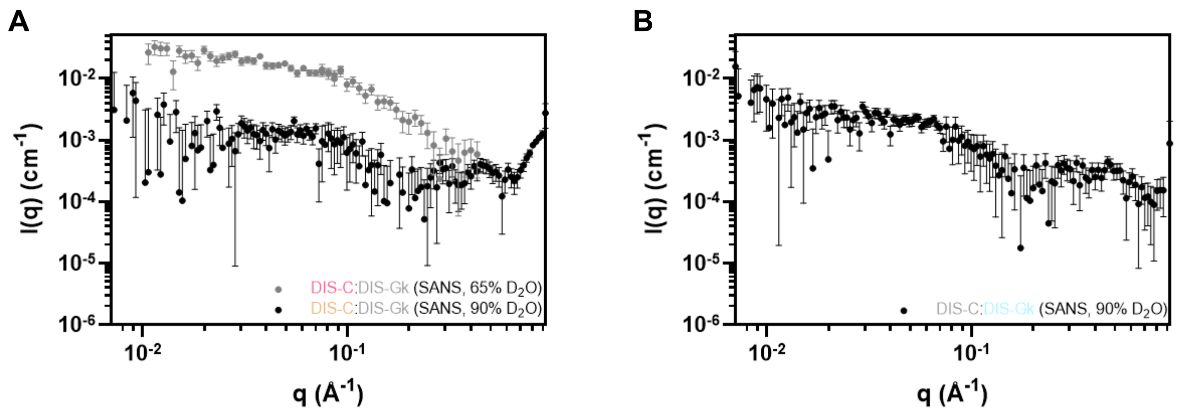


**Figure S4. Full experimental SANS scattering data. (A)** $^{2}$H-DIS-C:*$^{1}$H-DIS-Gk* in 65% $D_2O$ buffer (gray), $^{1}$H-DIS-C:*$^{2}$H(42%)-DIS-Gk* in 90% $D_2O$ buffer (black), and **(B)** *$^{2}$H(42%)-DIS-C*:$^{1}$H-DIS-Gk in 90% $D_2O$ buffer (black). Matched out components are italicized.

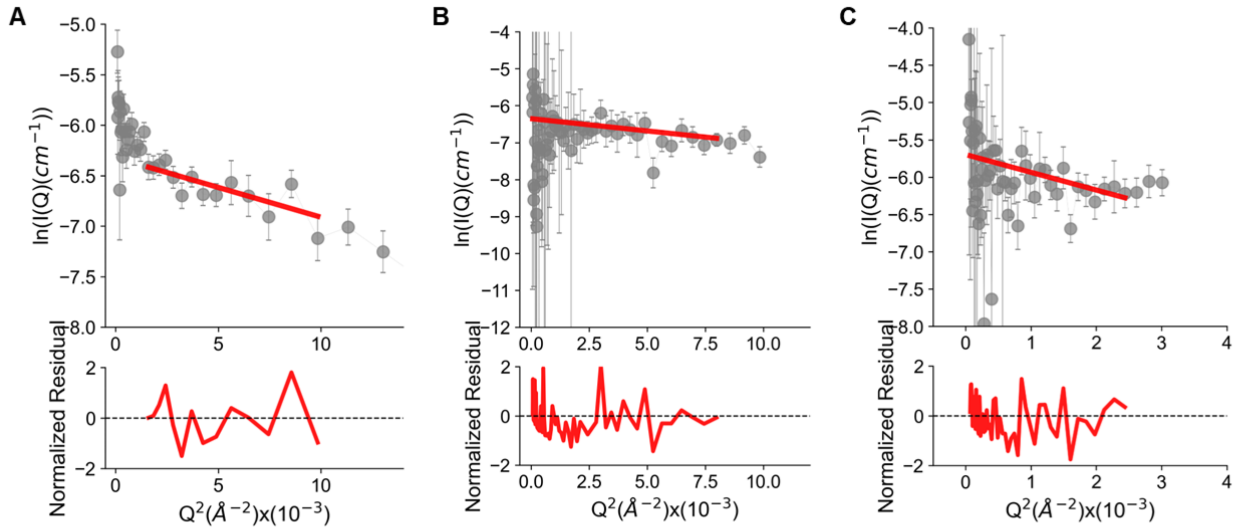


**Figure S5. SANS Guinier analysis of selectively deuterated DIS-C:DIS-Gk complexes.** Guinier fits (red) to experimental data (gray) for **(A)** $^{2}$H-DIS-C:*$^{1}$H-DIS-Gk* in 65% $D_2O$ buffer, **(B)** $^{1}$H-DIS-C:*$^{2}$H(42%)-DIS-Gk* in 90% $D_2O$ buffer, and **(C)** *$^{2}$H(42%)-DIS-C*:$^{1}$H-DIS-Gk in 90% $D_2O$ buffer. Matched out components are italicized.

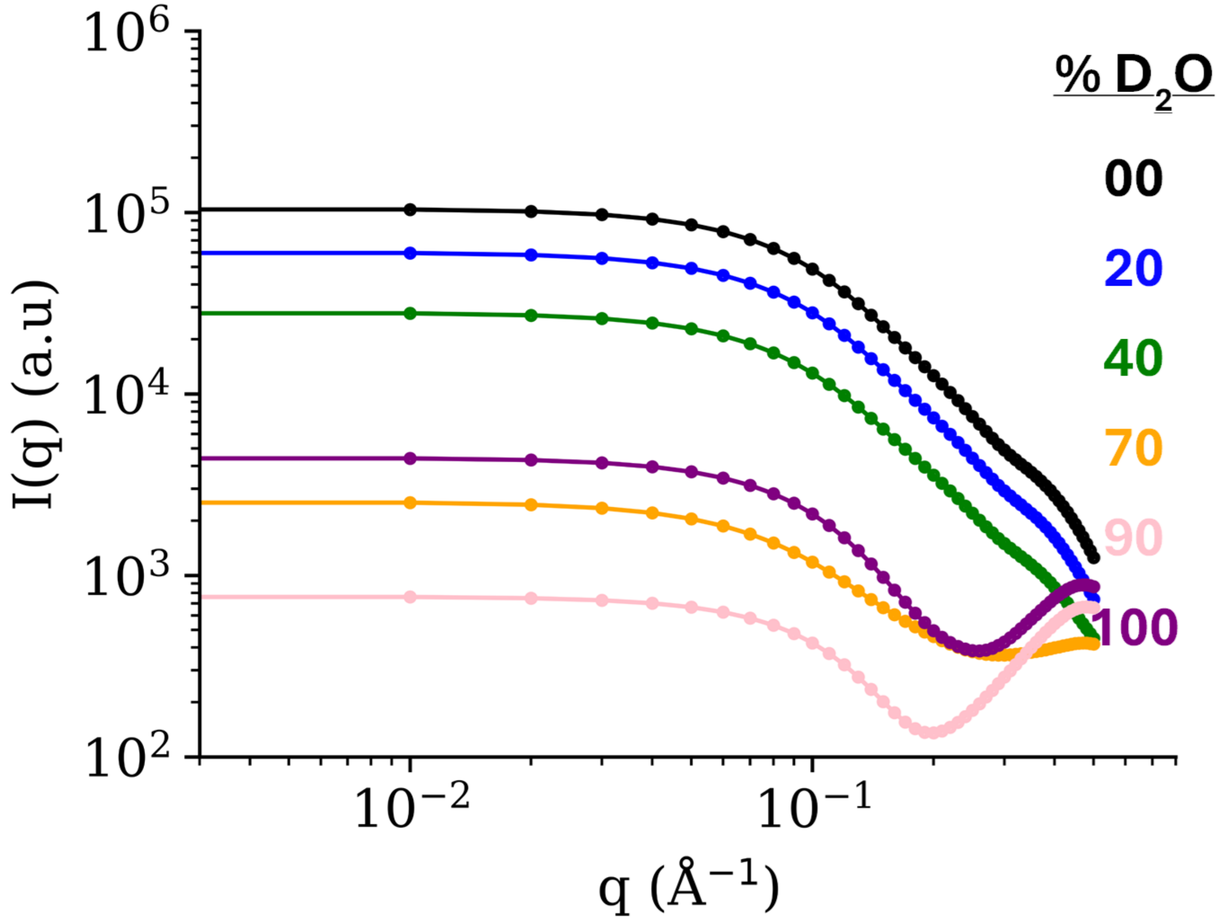


**Figure S6. Simulated SANS profiles of DIS-C at different contrast conditions show differences in the scattering data.** At low neutron contrast and smaller scattering vectors, the forward intensity increases, providing more information about the size and shape of the RNA. In contrast, at high neutron contrast and larger scattering vectors, the data reveals more details about the internal structure.

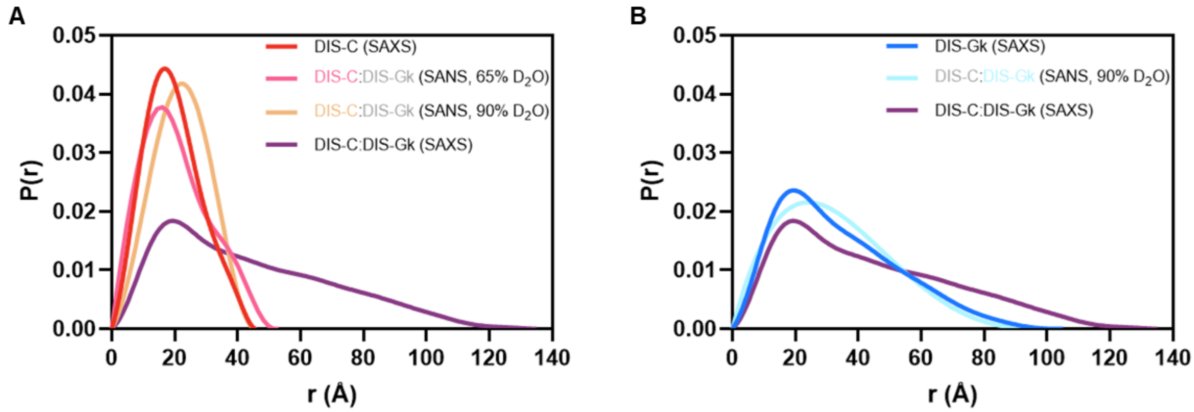


**Figure S7. $P(r)$ functions of the selectively deuterated DIS-C:DIS-Gk complexes show successful contrast matching.** Comparison of normalized pair distance distribution functions of **(A)** DIS-C and **(B)** DIS-Gk to the DIS-C:DIS-Gk complex indicates successful contrast matching. Matched out components are indicated in gray in the legend.

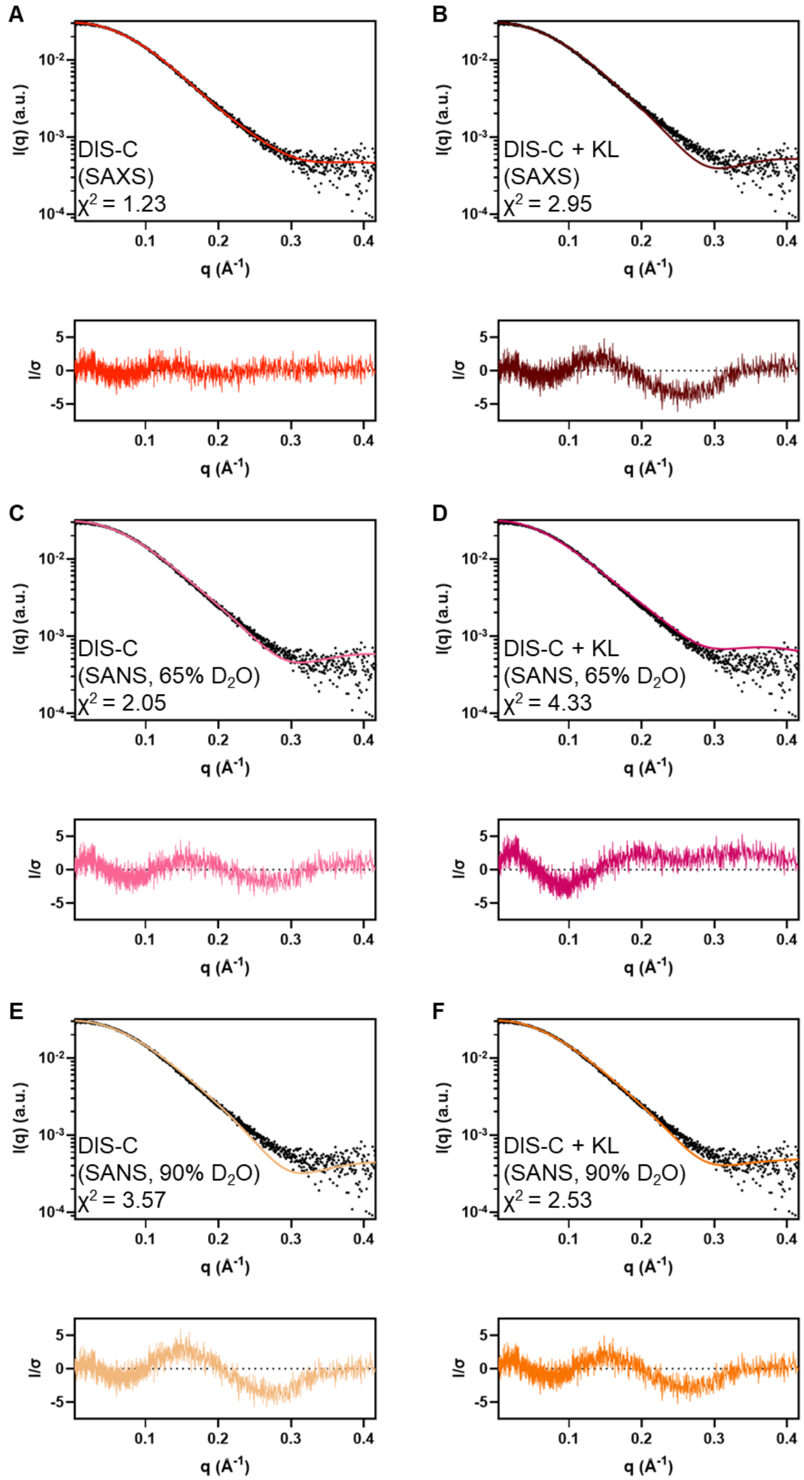


**Figure S8. Fits of final DIS-C models to DIS-C SAXS data.** Theoretical scattering (colored lines) of **(A)** DIS-C (SAXS), **(B)** DIS-C + KL (SAXS), **(C)** DIS-C (SANS, 65% $D_2O$), **(D)** DIS-C + KL (SANS, 65% $D_2O$), **(E)** DIS-C (SANS, 90% $D_2O$), and **(F)** DIS-C + KL (SANS, 90% $D_2O$) models fit to the DIS-C SAXS data (black dots) as reported in Table S3. Residuals are shown below each plot.

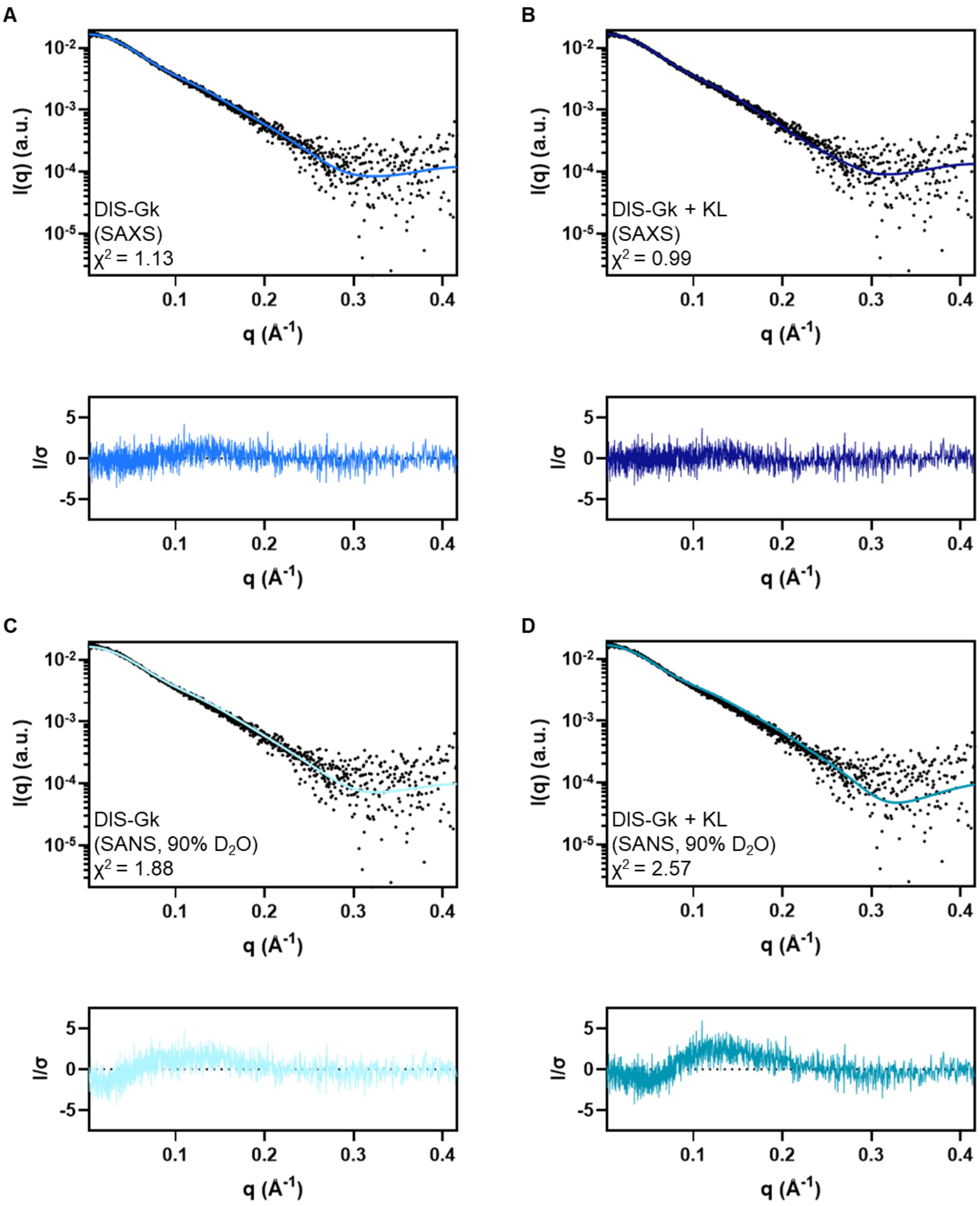


**Figure S9. Fits of final DIS-Gk models to DIS-Gk SAXS data.** Theoretical scattering (colored lines) of **(A)** DIS-Gk (SAXS), **(B)** DIS-Gk + KL (SAXS), **(C)** DIS-Gk (SANS, 90% $D_2O$), and **(D)** DIS-Gk + KL (SANS, 90% $D_2O$) models fit to the DIS-Gk SAXS data (black dots) as reported in Table S4. Residuals are shown below each plot.

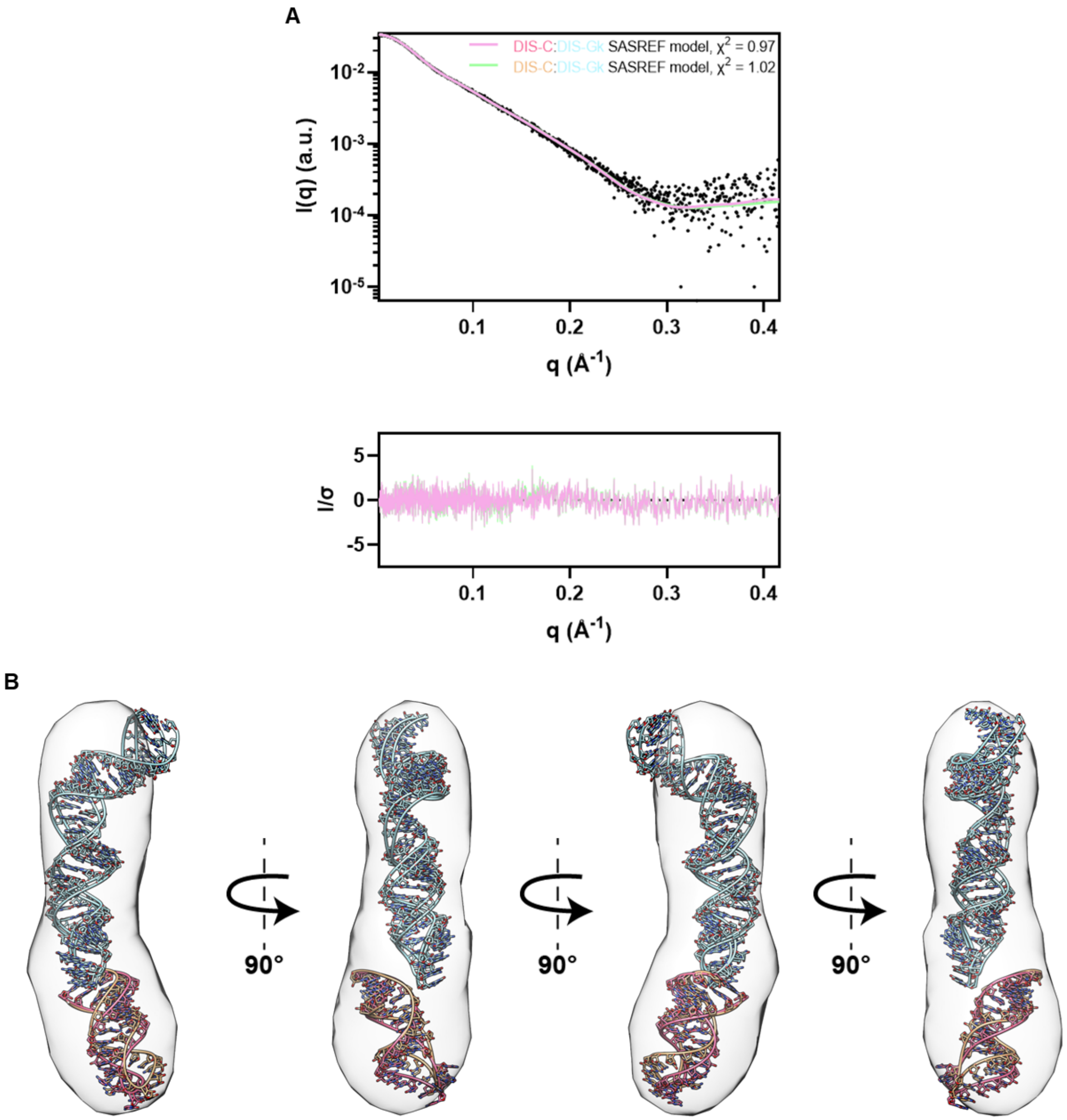


**Figure S10. Rigid body modeling of SANS-guided structures. (A)** SASREF rigid body modeling of structures in Figure 4 optimized to the DIS-C:DIS-Gk SAXS data fitted to the experimental SAXS data. DIS-C (SANS, 65% $D_2O$):DIS-Gk (SANS, 90% $D_2O$) is in light pink and DIS-C (SANS, 90% $D_2O$):DIS-Gk (SANS, 90% $D_2O$) is in light green. **(B)** SASREF rigid body modeling of structures in Figure 4 optimized to the DIS-C:DIS-Gk SAXS data fitted to the density map reconstruction (contour level: 7.5σ) from the DIS-C:DIS-Gk SAXS data. DIS-C SANS models are colored pink (65% $D_2O$) and light orange (90% $D_2O$), and the DIS-Gk SANS model is colored light teal (90% $D_2O$).

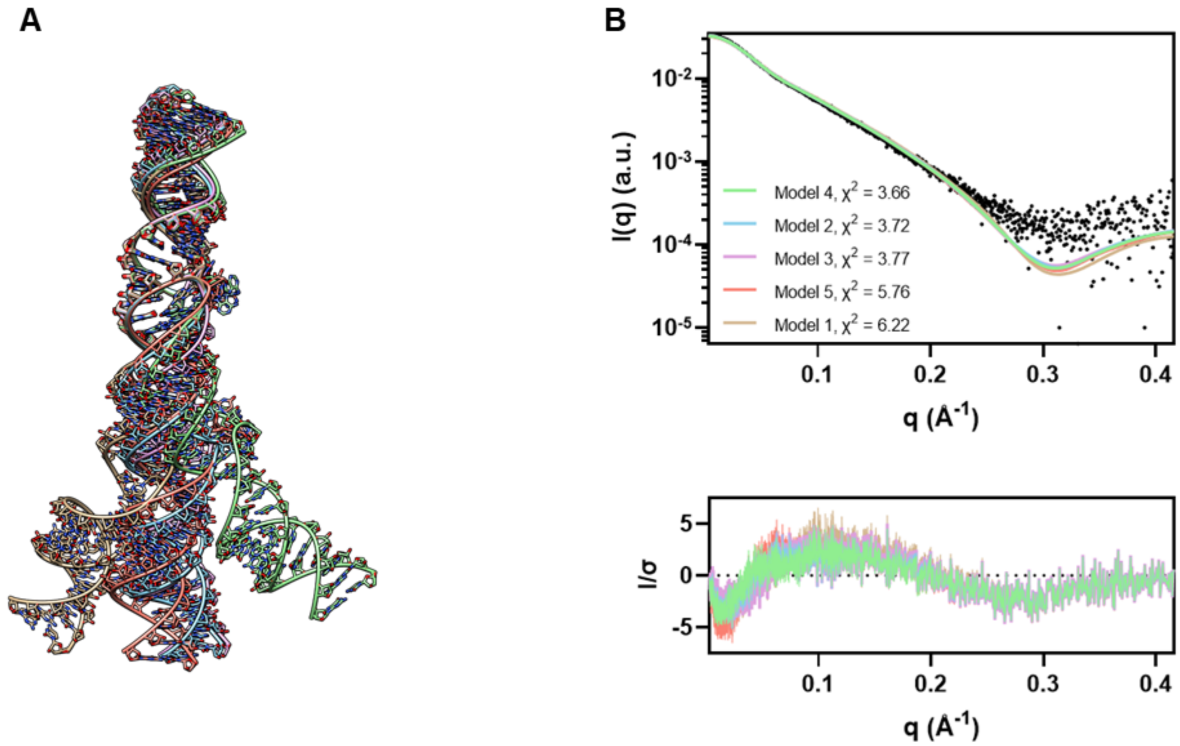


**Figure S11. AF3 DIS-C:DIS-Gk kissing complexes fit to experimental SAXS data. (A)** AF3 DIS-C:DIS-Gk predicted models 1 (tan), 2 (blue), 3 (pink), 4 (green), and 5 (red) aligned to model 4 at the kissing loop interface. **(B)** Theoretical scattering profiles of the AF3 models (colored lines) fit to the experimental DIS-C:DIS-Gk SAXS data (black dots, top) and corresponding residuals (bottom).

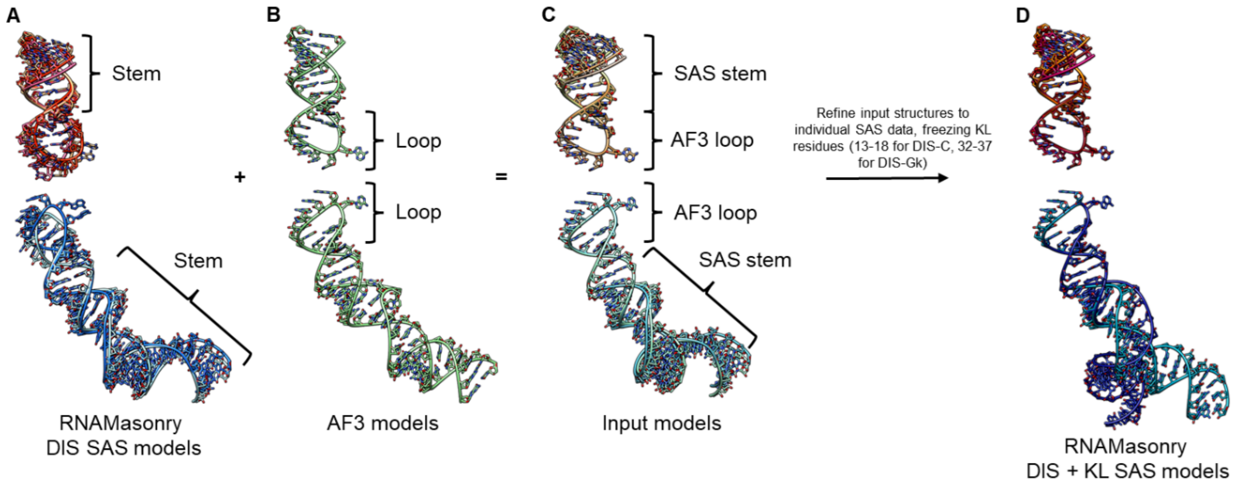


**Figure S12. Modeling of the DIS RNAs with kissing loops. (A)** The RNAMasonry DIS SAS models [DIS-C (top) and DIS-Gk (bottom)] reported in Figures 2 and 4 with the correct secondary structures. **(B)** AF3 model 4 of the DIS-C:DIS-Gk kissing complex, with the correct KL geometry but incorrect secondary structures. **(C)** Input structures for RNAMasonry generated by appending the stems of **(A)** and the apical loops of **(B)**. **(D)** RNAMasonry DIS + KL SAS models reported in Figure 5 where the input structures in **(C)** were further refined to their respective SAS data, keeping the KL residues frozen.

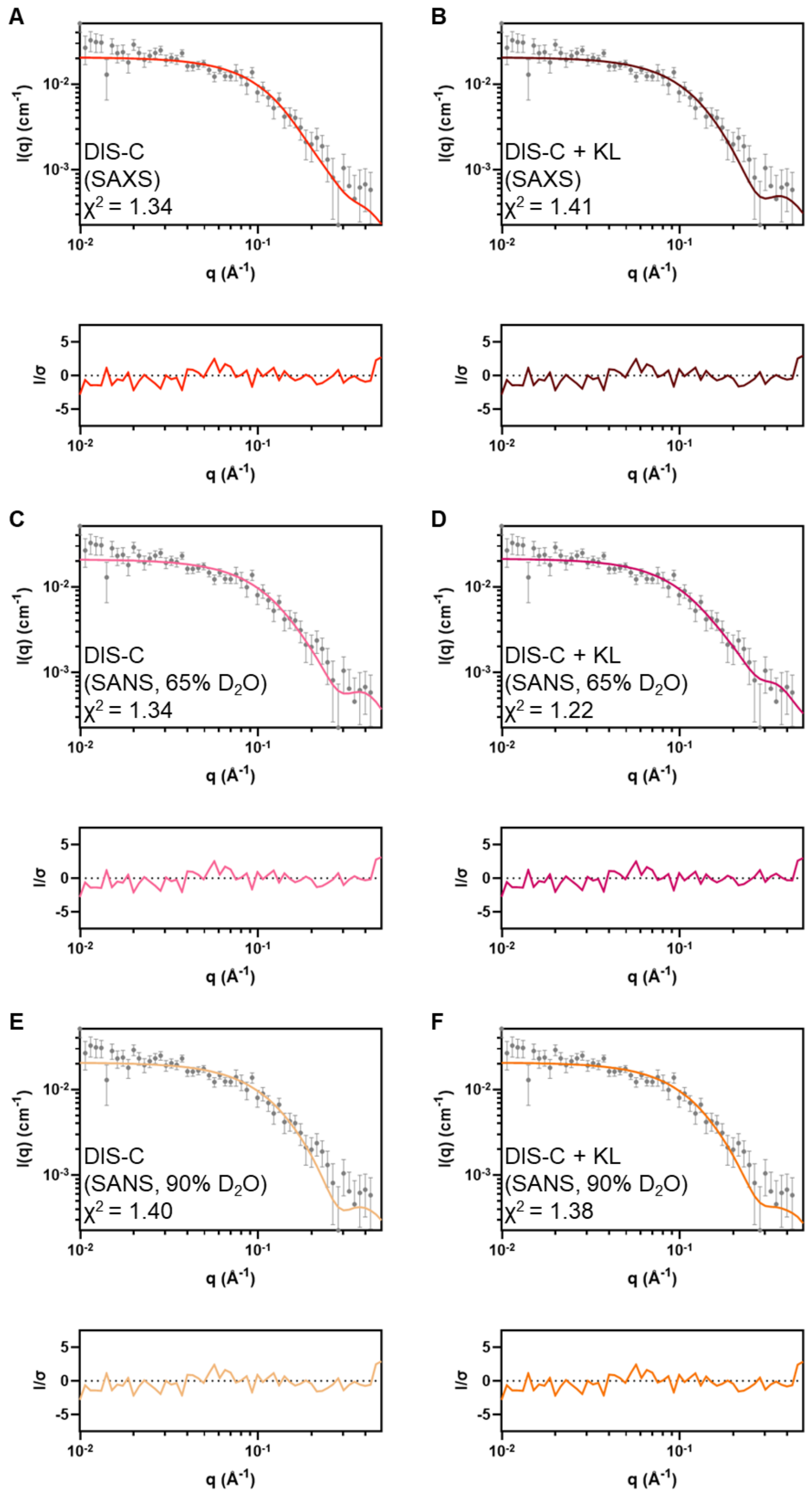


**Figure S13. Fits of final DIS-C models to $^2$H-DIS-C:$^1$H-DIS-Gk SANS (65% $D_2O$).** Theoretical scattering (colored lines) of **(A)** DIS-C (SAXS), **(B)** DIS-C + KL (SAXS), **(C)** DIS-C (SANS, 65% $D_2O$), **(D)** DIS-C + KL (SANS, 65% $D_2O$), **(E)** DIS-C (SANS, 90% $D_2O$), and **(F)** DIS-C + KL (SANS, 90% $D_2O$) models fit to the $^2$H-DIS-C:$^1$H-DIS-Gk (65% $D_2O$) SANS data (gray dots) as reported in Table S3. Residuals are shown below each plot.

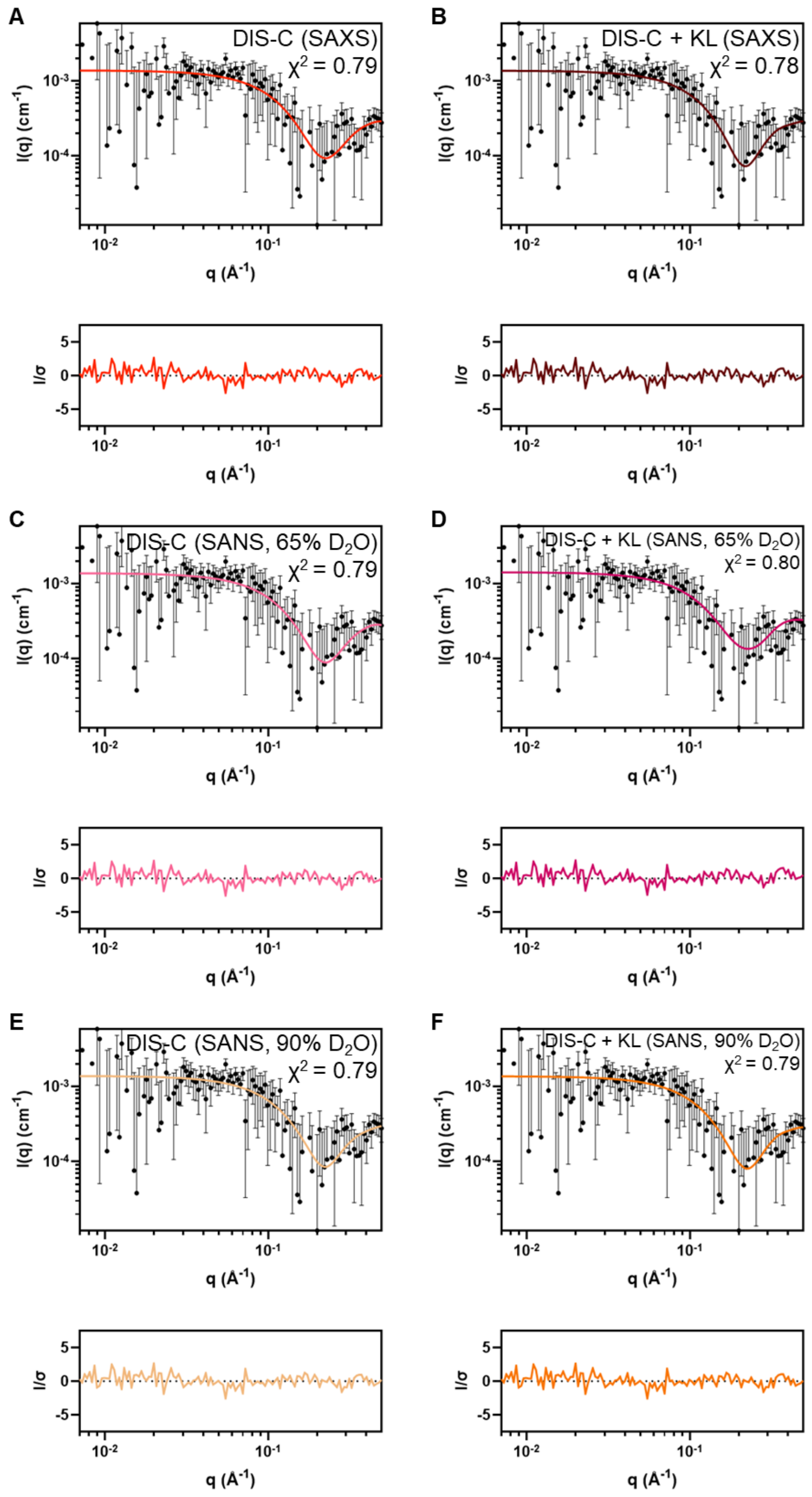


**Figure S14. Fits of final DIS-C models to $^{1}$H-DIS-C:$^{2}$H(42%)-DIS-Gk SANS (90% $D_2O$).** Theoretical scattering (colored lines) of **(A)** DIS-C (SAXS), **(B)** DIS-C + KL (SAXS), **(C)** DIS-C (SANS, 65% $D_2O$), **(D)** DIS-C + KL (SANS, 65% $D_2O$), **(E)** DIS-C (SANS, 90% $D_2O$), and **(F)** DIS-C + KL (SANS, 90% $D_2O$) models fit to the $^{1}$H-DIS-C:$^{2}$H(42%)-DIS-Gk (90% $D_2O$) SANS data (black dots) as reported in Table S3. Residuals are shown below each plot.

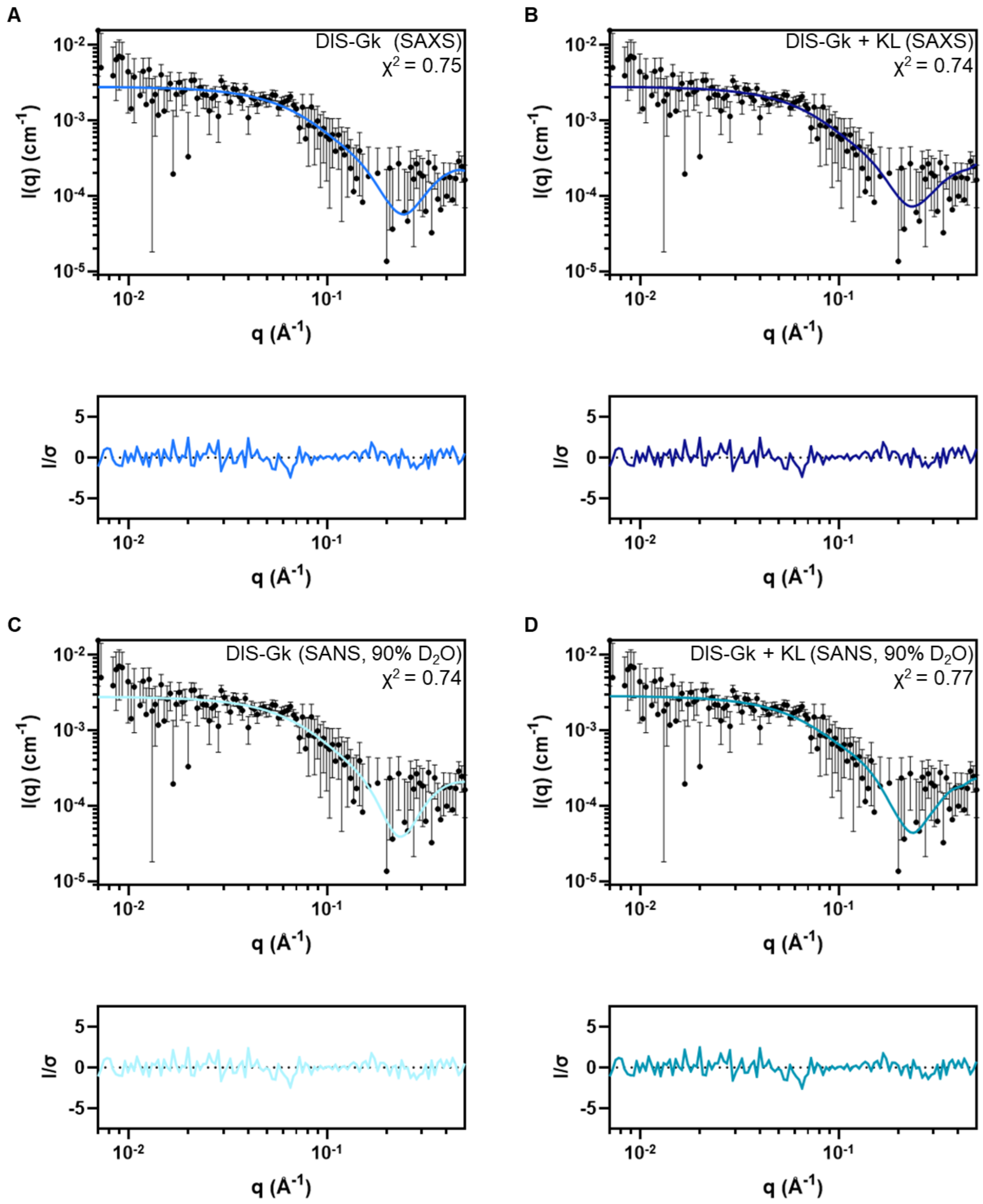


**Figure S15. Fits of final DIS-Gk models to $^{2}$H(42%)-DIS-C:$^{1}$H-DIS-Gk SANS (90% $D_2O$).** Theoretical scattering (colored lines) of **(A)** DIS-Gk (SAXS), **(B)** DIS-Gk + KL (SAXS), **(C)** DIS-Gk (SANS, 90% $D_2O$), and **(D)** DIS-Gk + KL (SANS, 90% $D_2O$) models to the $^{2}$H(42%)-DIS-C:$^{1}$H-DIS-Gk (90% $D_2O$) SANS data (black dots) as reported in Table S4. Residuals are shown below each plot.

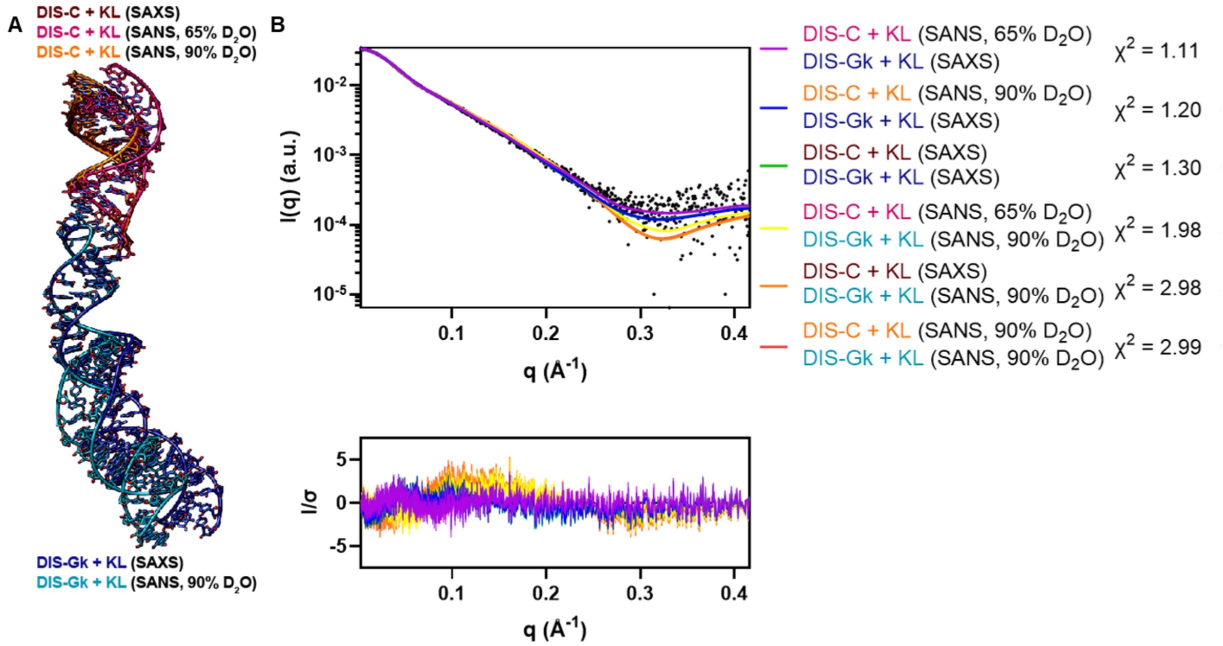


**Figure S16. SAS-guided DIS-C:DIS-Gk kissing complexes fit to experimental SAXS data. (A)** All combinations of the DIS-C:DIS-Gk kissing complexes composed of the individual DIS-C and DIS-Gk RNAs refined to their respective SAS data while keeping the stacking geometry of the KL residues predicted by AF3. The KL residues of the individual SAS-guided DIS RNAs were aligned to their respective RNAs KL residues in AF3 model 4. **(B)** Theoretical scattering profiles (colored lines) of the possible DIS-C:DIS-Gk complex models in **(A)** fit to the experimental DIS-C:DIS-Gk SAXS data (black dots, top) and corresponding residuals (bottom).

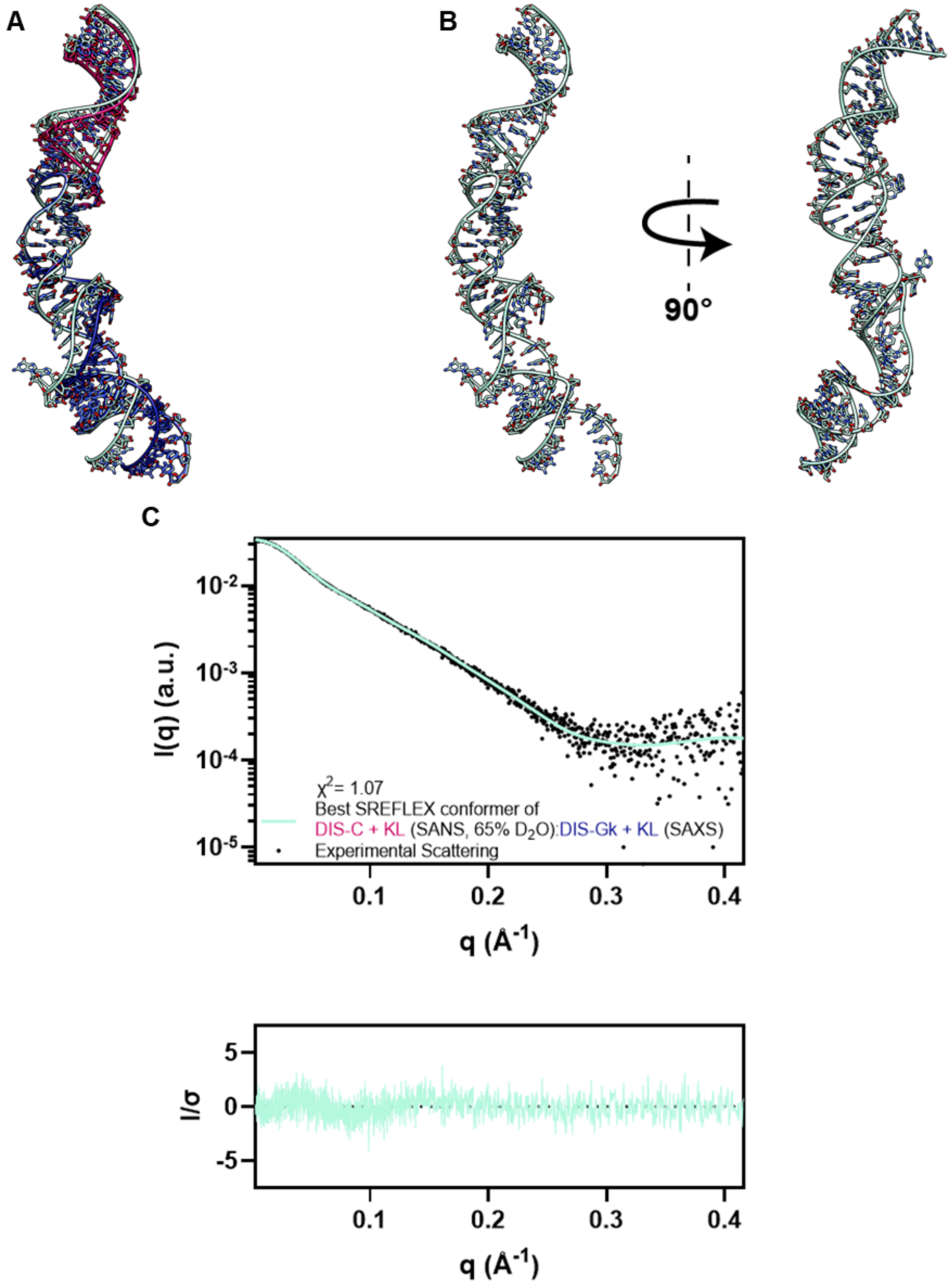


**Figure S17. Estimated flexibility of the DIS-C + KL (SANS, 65% $D_2O$):DIS-Gk + KL (SAXS) kissing complex. (A)** Alignment of the DIS-C + KL (SANS, 65% $D_2O$):DIS-Gk + KL (SAXS) kissing complex from Figure 5 to its best flexible conformer as estimated by SREFLEX (cyan), with an overall RMSD of 4.1 Å. **(B)** Structure of the best conformer optimized to the DIS-C:DIS-Gk SAXS data using the SAS-guided complex structure from Figure 5 as the input for SREFLEX. **(C)** Theoretical scattering of **(B)** fit to the DIS-C:DIS-Gk SAXS data.

Coordinate files (.pdb) of input and final structural models to support Figures 2, 4, 5, and S11 are uploaded.

Filenames:
S1 - DIS-C (SAXS).pdb
S2 - DIS-C + KL (SAXS) RNAMasonry input.pdb
S3 - DIS-C + KL (SAXS).pdb
S4 - DIS-C (SANS, 65d) pdb
S5 - DIS-C + KL (SANS, 65d) RNAMasonry input.pdb
S6 - DIS-C + KL (SANS, 65d).pdb
S7 - DIS-C (SANS, 90d).pdb
S8 - DIS-C + KL (SANS, 90d) RNAMasonry input.pdb
S9 - DIS-C + KL (SANS, 90d).pdb
S10 - DIS-Gk (SAXS).pdb
S11 - DIS-Gk + KL (SAXS) RNAMasonry input.pdb
S12 - DIS-Gk + KL (SAXS).pdb
S13 - DIS-Gk (SANS, 90d).pdb
S14 - DIS-Gk + KL (SANS, 90d) RNAMasonry input.pdb
S15 - DIS-Gk + KL (SANS, 90d).pdb
S16 - DIS-C + KL (SANS, 65d)_DIS-Gk + KL (SAXS) complex.pdb
